# SWARM resolves nanopore signal interference between RNA modification types and reveals splicing-shaped pseudouridylation

**DOI:** 10.64898/2025.12.18.695332

**Authors:** Stefan Prodic, Alice Cleynen, Shafi Mahmud, Akanksha Srivastava, Agin Ravindran, Madhu Kanchi, Arash Hajizadeh Dastjerdi, Aditya J. Sethi, Miona Ćorović, Ronit Jain, Marco Guarnacci, Gabriela Santos-Rodriguez, Grazi Vieira, Thomas Preiss, Robert J. Weatheritt, Rippei Hayashi, Gaetan Burgio, Nicole M. Martinez, Nikolay E. Shirokikh, Eduardo Eyras

**Affiliations:** EMBL Australia Partner Laboratory Network at the Australian National University, Canberra, ACT 2601, Australia; The Shine-Dalgarno Centre for RNA Innovation, The John Curtin School of Medical Research, Australian National University, Canberra, ACT 2601, Australia; The Centre for Computational Biomedical Sciences, The John Curtin School of Medical Research, Australian National University, Canberra, ACT 2601, Australia; France-Australia Mathematical Sciences and Interactions ANU - CNRS International Research Laboratory, Canberra, Australia; Institute of Molecular Biology (IMB), 55128 Mainz, Germany; Department of Chemical and Systems Biology, Stanford University, Stanford, CA 94305, USA; School of Biotechnology and Biomolecular Sciences, University of New South Wales, 2052 Sydney, Australia; EMBL Australia Partner Laboratory Network at the Garvan Institute of Medical Research, 2010 Sydney, Australia; Department of Developmental Biology, Stanford University, Stanford, CA 94305, USA; Sarafan ChEM-H Institute, Stanford University, Stanford, CA 94305, USA; Chan Zuckerberg Biohub, San Francisco, CA 94158, USA; Australian Centre for RNA Therapeutics in Cancer, School of Human Sciences, The University of Western Australia, Perth, WA

**Author notes:** These authors contributed equally (Alice Cleynen, Shafi Mahmud, Akanksha Srivastava).

## Abstract

Nanopore direct RNA sequencing promises to decode the epitranscriptome by detecting multiple modifications on individual RNA molecules, but its potential for biological discovery is hampered by high false-positive rates. We present SWARM, an AI-based framework designed to overcome this fundamental limitation. Its key innovation is a crosstalk-aware training strategy that incorporates non-target modifications and orthogonally validated cellular signals, enabling high-precision detection of m6A, pseudouridine (Ψ), and m5C at single-nucleotide and single-molecule resolution. Using rigorous *in vitro* and cellular RNA benchmarks, SWARM outperforms existing tools and maintains strong agreement with orthogonal methods. Applying SWARM across mammalian tissues reveals thousands of novel modification sites with confirmed motifs and localisation patterns. Our high-resolution multi-tissue modification map revealed no evidence of widespread m6A-Ψ interplay in predominant writer contexts, challenging models of a coordinated epitranscriptomic code. We further discovered a previously unrecognised splicing-shaped mode of Ψ deposition, whereby TRUB1-mediated pseudouridylation preferentially occurs after exon-exon ligation, consistent with local RNA structure stabilisation. SWARM provides a robust, universally applicable tool for epitranscriptome discovery.

## Introduction

Recent years have seen a remarkable expansion in our understanding of RNA ^1–3^. In addition to the four canonical ribonucleotides, which make up the main backbone of RNA, there are over 170 epitranscriptomic modifications that potentially provide an extra dimension of RNA processing and control ^1–3^. Multiple research efforts have demonstrated the ability of modifications to modulate the expression and fate of diverse RNA types, including messenger RNA (mRNA), through the interplay with structure, protein-RNA interactions, stability, splicing, translation efficiency, and subcellular localisation ^4,5^.

The current evidence highlights two main characteristics. First, RNA modifications can be dynamic, with modified sites changing across specific conditions, such as developmental stages, cell types, and disease states ^1,6^. Second, the position in the mRNA appears to be functionally relevant, with modified sites being evolutionarily conserved and likely triggering positional-dependent outcomes ^7,8^. However, the precise mechanisms by which RNA modifications affect mRNA processing and function are not yet fully understood ^9,10^, largely because most studies are limited to one modification type or one mRNA molecule at a time.

It is thus essential to develop methods that accurately identify multiple modifications at single-molecule and single-nucleotide resolution that are easily applicable across multiple conditions. While several experimental methods exist, they usually detect one single modification at a time, require a non-modified control, and involve complex protocols, including RNA immunoprecipitation ^11^, chemical conversion ^12^, or enzymatic treatment ^13^.

Direct RNA sequencing (DRS) using molecular nanopores offers the potential to identify multiple modifications within an individual RNA molecule at a transcriptome-wide scale ^14–18^. However, a fundamental unresolved issue is signal crosstalk, where the presence of one modification alters the nanopore signal of a neighbouring nucleotide, leading to pervasive false positives in multi-modification detection, and confounding biological interpretation ^14–18^. Furthermore, currently published nanopore-based methods predominantly rely on training with synthetic signals, either from *in vitro* transcribed (IVT) synthetic RNAs with unnaturally dense modification contexts ^15,16^ or from synthetic oligos with limited sequence contexts ^19,20^. While there are methods that use signals from cellular mRNA ^14,21,22^ or rRNA ^14^, they are affected by a high proportion of low-stoichiometry sites that induce noise during training. Another significant limitation of current nanopore-based methods for modification detection is a lack of clarity on the optimal parameters to use in single-sample detection, with common use of simple stoichiometry cutoffs. While this may work well for orthogonal experimental methods with negligible false-positive rates, it results in low precision and a high level of noise in nanopore predictions, especially for rare modifications, precluding the identification of biologically sound results.

To address these challenges, we developed SWARM (Single-molecule Workflow for Analysing RNA Modifications), an AI-based computational framework for the transcriptome-wide detection of N6-methyladenosine (m6A), 5-methylcytosine (m5C), and pseudouridine (Ψ) in both nanopore chemistries (SQK-RNA002 and SQK-RNA004) at single-nucleotide resolution in individual molecules from a single sample without the need for controls. We introduce three key innovations with respect to current strategies ^23^: (1) Incorporation of non-target modifications in training data to minimise signal crosstalk, effectively reducing false positives; (2) diverse training data including cell-line signals from orthogonally validated sites on mRNA, rRNA, and tRNA, enabling robust calling of modified nucleotides and accurate stoichiometry estimates in cellular RNA; (3) systematic model selection and benchmarking using synthetic and cellular RNA, ensuring specificity for the target modification, as well as high precision for low-frequency target modifications.

These innovations led to two novel biological insights. A comprehensive analysis of multiple tissues across five mammalian species, as well as perturbation experiments, revealed that, contrary to current models, there is no evidence of widespread coordination between m6A and Ψ in their predominant writer contexts. Moreover, through our comprehensive analyses, we discovered a novel mode of Ψ deposition shaped by splicing, in which TRUB1-mediated modification near splice sites preferentially occurs after the right exon-exon pair is expressed, revealing a new link between mRNA maturation and epitranscriptomic modification.

## Results

### Refined training workflow underpins accurate analysis of multiple RNA modifications

SWARM identifies N6-methyladenosine (m6A), 5-methylcytosine (m5C), and pseudouridine (Ψ), with single-base and single-molecule resolution, testing every A/C/U base at any sequence context on every read, using as input the nanopore signal aligned to a 9-mer sequence centred on the position of interest (Fig. 1a). SWARM read-level models were trained using mixtures of nanopore signals from *in vitro* transcribed (IVT) synthetic RNA templates (Supp. Table 1), supplemented with nanopore signals from sites in cellular RNA validated by orthogonal experiments (Supp. Table 2), thereby expanding the modification contexts. These cellular sites encompassed mRNA m6A sites validated by GLORI ^12^, rRNA Ψ sites validated by mass spectrometry and BACS ^24,25^, and tRNA m5C sites validated by UBS ^26^ (Fig. 1a). Specifically, positive signals were taken from validated high-stoichiometry sites from direct RNA sequencing in cell types matching the orthogonal experiment samples, while negative signals were obtained from non-modified IVT cellular and synthetic RNA, with genomic coordinates and sequence contexts matching the positive data (see Methods for details). To account for the differences in training data composition, we considered three neural network architectures with varying parameter counts, referred to as Mini, Mid, and Large (Methods), for each modification type and assessed their suitability across different benchmarks.

**Figure 1.**
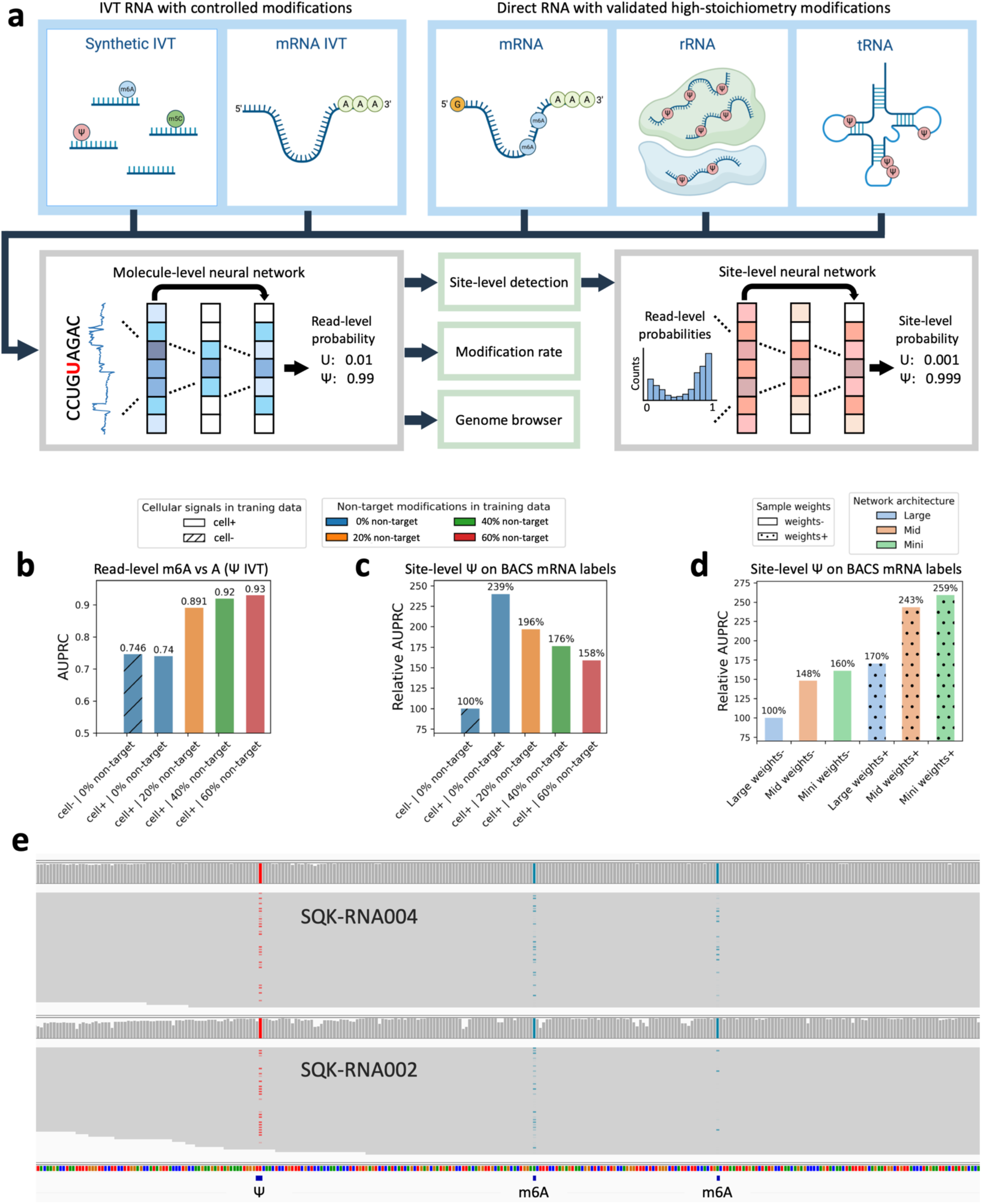
SWARM framework for analysing multiple RNA modifications. **(a)** SWARM’s workflow facilitates the training of read-level RNA modification models with synthetic and cell-derived nanopore signals to predict modifications in individual molecules at single nucleotide resolution. The read-level models are the baseline for modification visualisation in a genome browser, measurement of modification stoichiometry and detection of modification sites. Site-level detection of modifications is achieved with a downstream neural network predicting modification states at a reference site coordinate from the distribution of read-level probabilities. **(b-c)** Bar plots indicating impact of training data composition on model performance, with % of non-target modifications in negatives shown in colours, and inclusion of cellular data during training indicated with hatches. Bar plot (b) showing area under the precision-recall curve (AUPRC) for the SWARM m6A SQK-RNA002 models (Mini network) computed using read-level predictions on A sites from the m6A IVT as positives and A sites from the Ψ IVT as negatives. Bar plot (c) with relative AUPRC for the SWARM Ψ SQK-RNA002 models (Mini network) using site-level predictions on HeLa mRNA evaluated with BACS sites. Relative AUPRC was calculated with respect to the model with 0% non-target modifications and no cellular data. **(d)** Bar plot showing the impact of training hyperparameters (network architecture and sample weights) on the mRNA site-level detection. Relative AUPRC for the SWARM Ψ SQK-RNA002 models was computed using site-level predictions on HeLa mRNA evaluated with BACS sites, with values shown relative to the model with “Large” architecture and no sample weights. Model architecture is shown with colours, while usage of sample weights is indicated with dots. **(e)** IGV snapshot of SWARM outputs showing per-read single-base probabilities for a Ψ site (red, position 3567), which was also detected by BID-seq in HeLa ^30^. In the same transcript, we show SWARM m6A single-molecule predictions (blue) with sites detected at positions 3654 and 3711, which were also reported by GLORI in HeLa ^12^. Validated sites are indicated under the sequence track and were converted from genomic to transcriptomic positions using the Ensembl hg38 GTF annotation.

**Table 1.** Single-read precision/recall analysis. The table displays the area under the precision-recall curves (AUPRC) for the benchmarking of different read-level modification tools for both nanopore chemistries (SQK-RNA002 and SQK-RNA004) at single-nucleotide resolution. Benchmarking datasets were built from independent synthetic IVTs and sequence contexts not used for training. Positive cases were built from reads containing the modification indicated in the column “Target modification”. Various negative sets were considered (“negatives”), encompassing reads without modifications (“Unmodified”) or reads containing non-target modifications; n.a.= not applicable.

| Target Modification | ONT kit | Tool | Negatives for the Area Under the Precision-Recall Curve (AUPRC) |  |  |  |
| --- | --- | --- | --- | --- | --- | --- |
| | | | Unmodified | $\Psi$ | m5C | m6A |
| m6A | SQK-RNA002 | SWARM | **0.95** | **0.9** | **0.91** | n.a. |
|  |  | mAFiA | 0.92 | 0.81 | 0.80 | n.a. |
|  |  | TandemMod | 0.90 | 0.70 | 0.74 | n.a. |
|  |  | SingleMod | 0.86 | 0.82 | 0.76 | n.a. |
|  |  | CHEUI | 0.86 | 0.73 | 0.78 | n.a. |
|  |  | m6anet | 0.82 | 0.78 | 0.73 | n.a. |
|  |  | m6ABC | 0.74 | 0.73 | 0.62 | n.a. |
|  | SQK-RNA004 | SWARM | **0.96** | **0.90** | **0.92** | n.a. |
|  |  | dorado | 0.94 | 0.88 | 0.89 | n.a. |
|  |  | m6Anet | 0.78 | 0.75 | 0.66 | n.a. |
|  |  | SingleMod | 0.65 | 0.63 | 0.64 | n.a. |
| $\Psi$ | SQK-RNA002 | SWARM | **0.97** | n.a. | **0.92** | **0.92** |
|  |  | psi-co-mAFiA | 0.88 | n.a. | 0.80 | 0.85 |
|  |  | TandemMod | 0.80 | n.a. | 0.57 | 0.69 |
|  | SQK-RNA004 | SWARM | **0.98** | n.a. | **0.92** | **0.93** |
|  |  | dorado | 0.93 | n.a. | 0.88 | 0.90 |
| m5C | SQK-RNA002 | SWARM | **0.96** | **0.89** | n.a. | **0.93** |
|  |  | TandemMod | 0.96 | 0.87 | n.a. | 0.88 |
|  |  | CHEUI | 0.91 | 0.78 | n.a. | 0.87 |
|  | SQK-RNA004 | SWARM | **0.99** | **0.94** | n.a. | **0.96** |
|  |  | dorado | 0.88 | 0.80 | n.a. | 0.84 |

One key component of our training strategy was designed to address the elevated false-positive rate in nanopore RNA modification prediction caused by non-target RNA modifications ^16,27^. We supplemented the negative read-level training data with IVT nanopore signals containing non-target modifications. We then tested the accuracy of the three network architectures, each trained on a different set of proportions (0%, 20%, 40%, 60%) of non-target modifications in the negative training data examples. This resulted in the drastic improvement of the separation of target from non-target modification in nanopore signals not seen during training (Fig. 1b) (Supp. Fig. 1). The improvement was most prominent for resolving the prediction errors caused by the m6A models when Ψ was present in the negative test set. Moreover, using 20% non-target modifications led to a larger jump in performance improvement, while 40% and 60% showed diminishing improvements (Fig. 1b) (Supp. Fig. 1). In contrast, inclusion of reads from cellular sites in the training data did not improve the accuracy of the read-level models (Fig. 1b) (Supp. Fig. 1).

An additional key aspect of our framework aimed to account for the low frequency of modifications in cellular mRNA ^28,29^. This makes the transcriptome-wide detection of modifications challenging, as it requires a false-positive rate below the actual modification frequency to yield meaningful results. To address this, we implemented a site-level neural network designed to estimate the level of confidence about the presence of a modification at a specific transcript reference position, taking as input the distribution of probabilities across individual reads mapped to that site (Fig. 1a). Site-level models were trained with data composed of synthetic combinations of IVT signals in assorted proportions of modified and unmodified reads spanning a wide range of stoichiometry and coverage levels, considering sites as modified if their stoichiometry was over 10% (Methods). To assess mRNA modification site detection across different read-level model hyperparameters, we trained a site-level model with hyperparameters consistent for each read-level model and evaluated the site-level predictions against orthogonally reported mRNA modification sites: m6A model hyperparameters were evaluated on GLORI sites in Hek293T cells, Ψ models on BACS sites in HeLa cells, and m5C on UBS sites in HeLa cells. These tests showed that inclusion of cellular signals in the training of the read-level models led to large improvements in accuracy of the site-level predictions compared to using only IVT signals (Fig. 1c) (Supp. Fig. 2). In contrast, using more than 20% of negatives with non-target modifications in the read-level model training had a negative impact on the accuracy of site-level detection (Fig. 1c) (Supp. Fig. 2). Based on these results, we selected 20% non-target modification in the negatives and cellular data inclusion as the training dataset for read-level models.

The final selection of hyperparameters involved choosing the optimal architecture among the three tested ones (Mini, Mid, Large) and the training sample weights between the IVT and cellular data (Supp. Table 1) (Methods) for each modification model and sequencing chemistry. To this end, we again aimed at balancing accuracy at the read and site levels. We observed that the “Large” architecture had the best performance on IVT read-level benchmarks for all modifications (Supp. Fig. 3), while the smaller “Mini” and “Mid” read-level architectures often generalised better for site-level detection (Fig. 1d) (Supp. Fig. 3). Moreover, usage of sample weights generally resulted in no improvements for read-level or site-level performance, except for the SQK-RNA002 Ψ model, where balanced weighting of IVT and cellular rRNA signals drastically improved mRNA site detection (Fig. 1d) (Supp. Fig. 3). We thus decided to proceed with a neural network architecture and sample weights configuration uniquely tailored for each modification model, selecting models with the highest mRNA site overlap with orthogonal methods (Supp. Fig. 3), except for the m6A model for SQK-RNA004, which was selected according to the best read-level perfromance, given the small differences in mRNA orthogonal overlap and the observed variability in performance on IVT controls (Supp. Fig. 3).

Each one of the SWARM models outputs high-resolution modification readouts in the standard modSAM format, enabling visualisation of modified reference sites and individual molecules in genome browsers (Fig. 1e). As an example, we illustrate the SWARM output for the human transcript ENST00000367876.9, with a Ψ site also detected by BID-seq ^30^ and two m6A sites also detected by GLORI ^12^ in different studies. The IGV snapshot highlights molecule-level detection of Ψ and m6A, enabling high-resolution visualization of epitranscriptomic marks and suggesting relationships between modifications within individual molecules.

### SWARM achieves unmatched modification specificity across synthetic controls

To compare SWARM models with other tools for the read-level detection of RNA modifications, we used nanopore signals from independent IVT experiments corresponding to IVT sequences not used in our training data and hyperparameter selection, using as positives individual signals containing the target modification and as negatives individual signals having either no modifications or non-target modifications. SWARM consistently maintains the highest separation of target and non-target modifications for m6A, Ψ, and m5C detection on both sequencing chemistries (Table 1) (Supp. Fig. 4). In contrast, the prediction accuracy of other methods decreases when applied to all the tested non-target modifications (Table 1) (Supp. Fig. 4).

In particular, Ψ had the strongest impact as a non-target modification affecting m6A and m5C prediction models (Table 1). Another notable insight is the drop in modification separation observed for the TandemMod Ψ model when introducing m5C in the negative test set (Table 1), with AUPRC = 0.57 when separating Ψ from Uridines in m5C background. This might be explained by the fact that TandemMod was fine-tuned on sparse Ψ data from the original m5C model. These results illustrate the importance of testing and reporting the effect of non-target modifications on nanopore modification prediction, especially in settings where a model for one modification is fine-tuned from another modification model.

### Modification stoichiometry in mRNA and rRNA correlates with orthogonal datasets

To evaluate read-level detection in cellular datasets, we applied SWARM to HeLa and HEK293T sequencing datasets and compared the stoichiometry, i.e. the modification rate of the aggregated reads at each transcriptomic site, with orthogonal experimental methods performed in the same cell lines. This provides an independent biological test, as SWARM model training and hyperparameter selection were conducted agnostic of stoichiometry estimation. For m6A, SWARM showed a high stoichiometry correlation with GLORI ^12^ in HeLa mRNA (R>0.9) in both ONT sequencing chemistries, demonstrating improvement over most of previously published methods (Figs. 2a & 2b) (Supp. Figs. 5a & 5b). Notably, SingleMod ^31^, trained on GLORI stoichiometry, and dorado showed higher correlations with GLORI for the SQK-RNA004 chemistry.

**Figure 2.**
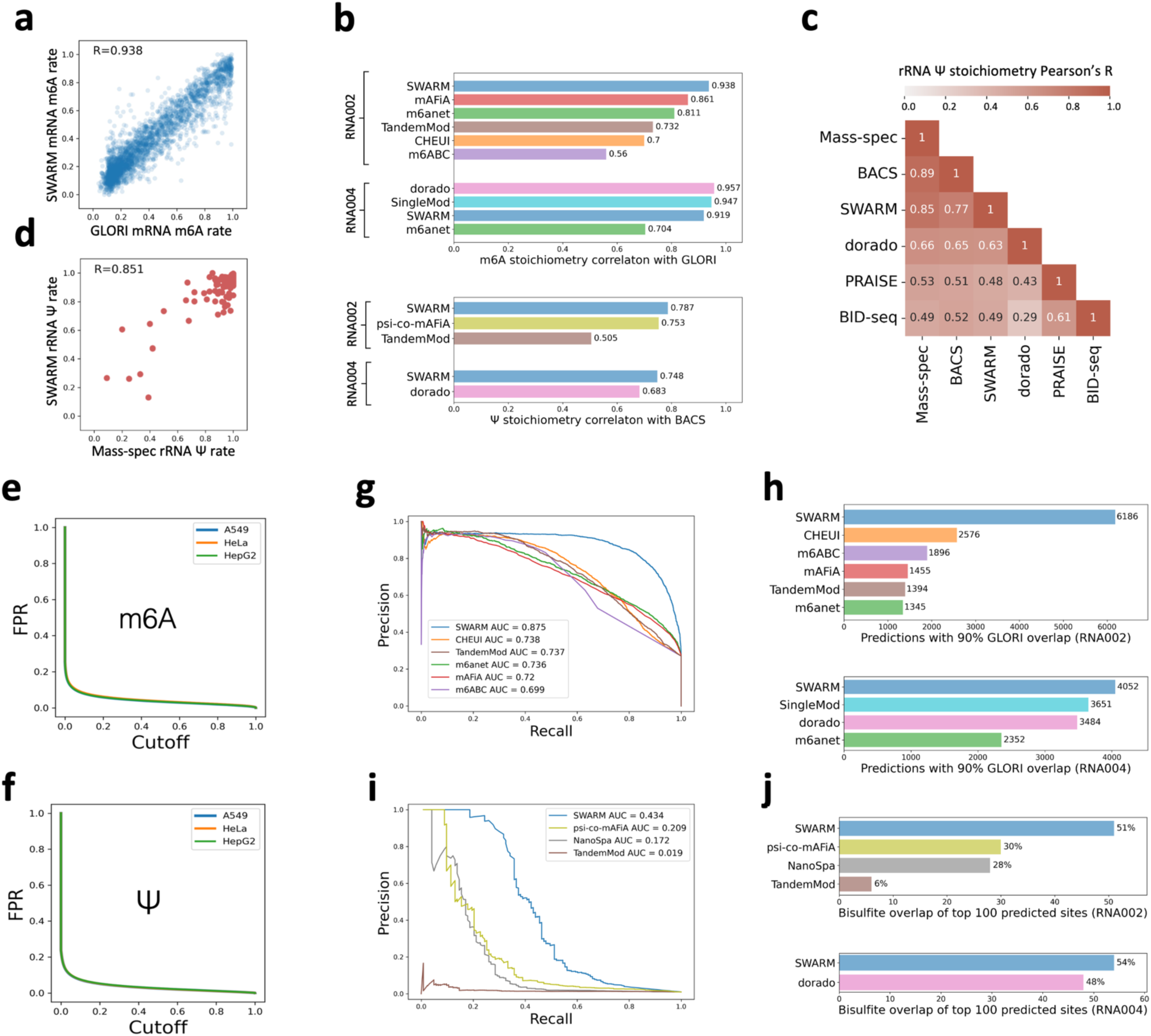
Stoichiometry and site-detection benchmarks on human cell lines. **(a)** Correlation of stoichiometry from GLORI (x-axis) and SWARM (y-axis) in HeLa SQK-RNA002 sample. **(b)** Pearson’s correlation coefficient (y-axis) for m6A (upper panel) and Ψ (lower panel) stoichiometries by each method (x-axis) for SQK-RNA002 and SQK-RNA004 chemistries. The correlations were based on the comparison with GLORI in HeLa for m6A, with BACS in HEK293T for Ψ. **(c)** Matrix of Pearson’s correlation coefficients for the pairwise comparison of Ψ stoichiometry reported by mass-spectrometry (MS), nanopore-based (SWARM, dorado), bisulfite-based (BID-seq, PRAISE), and BACS. **(d)** Correlation of Ψ stoichiometry in rRNA from mass spectrometry (x-axis) and SWARM (y-axis) in a HeLa SKQ-RNA004 sample. **(e-f)** SWARM’s False positive rate (FPR) (y-axis) as a function of the site-level probability cutoff (x-axis) for the m6A (e) and Ψ (f) models for SQK-RNA002 *in vitro* transcribed samples. **(g)** Precision-recall curve for SWARM and various other methods for the prediction of m6A at METTL3 candidate sites (DRACH) in SQK-RNA002, using as ground truth GLORI sites in HeLa. **(h)** Total number of transcriptomic DRACH sites (x-axis) predicted by each method (y-axis) in SQK-RNA002 (upper panel) and SQK-RNA004 (lower panel) at 90% precision, i.e. with 90% of the predictions validated by a GLORI. **(i)** Precision-recall curve for SWARM and other methods for the prediction of Ψ in PUS7 (UNΨAR, N=A,C,G,U) and TRUB1 (GUΨCNA) candidate sequence contexts for SQK-RNA002 DRS in HEK293T, using as benchmark the transcriptomic Ψ sites predicted by BID-seq or PRAISE in HEK293T. **(j)** For each method, we show the proportion of sites (x-axis) that overlap with the bisulfite methods BID-seq and PRAISE in HEK293T, considering the top 100 predicted sites by each method, ranked by the metric for site confidence given by the method, i.e. site-level probability for SWARM and NanoSPA, and stoichiometry for psi-co-mAFiA, TandemMod, and dorado. We show the results for SQK-RNA002 (upper panel) or SQK-RNA004 (lower panel). Only PUS7 (UNΨAR) and TRUB1 (GUΨCNA) sites were considered.

Regarding Ψ, our analyses in HEK293T mRNA showed that SWARM had the highest stoichiometry correlation with BACS ^25^ compared to other nanopore methods for both ONT chemistries (Fig. 2b) (Supp. Figs. 5c & 5d). To illustrate the level of agreement across different orthogonal methods for pseudouridine detection, we measured the Ψ stoichiometry with SWARM and dorado in a HeLa (SQK-RNA004) for ribosomal RNA (rRNA) and benchmarked it against the stoichiometry reported in HeLa by mass-spectrometry (MS) ^24^, considered to be closest to the ground truth, and multiple bisulfite-based methods ^25,30,32^ (Figs. 2c & 2d) (Supp. Fig. 6). BACS displayed the highest correlation with MS (R=0.89), followed closely by SWARM (R=0.85), while dorado and bisulfite-based methods displayed lower correlation values (R<0.7) (Fig. 2c) (Supp. Fig. 6). Similar to mRNA results, SWARM and BACS showed a high correlation with each other on rRNA (R=0.77).

In contrast to m6A and Ψ, currently available nanopore methods showed only a modest correlation of m5C stoichiometry with bisulfite-based techniques ^26^ in quantifying m5C in HEK293T mRNA (Supp. Figs. 5e & 5f). This discrepancy highlights a key limitation in the field, possibly due to inherent inaccuracies in existing detection methods or the challenges of reliably identifying m5C sites with low stoichiometry, a hallmark feature of this modification ^33^.

### SWARM bolsters precise transcriptome-wide detection of low-frequency modification sites

To evaluate cutoffs for the *de novo* detection of modified sites, we applied each site-level model on unmodified IVT transcriptomes from HeLa, HEK293T, and HepG2 ^34^ and estimated the false-positive rate across all 9-mer sequence contexts. The false positive rate values as a function of the site-level probability cutoffs remained similar across all three cell lines (A549, HeLa, HepG2) (Figs. 2e & 2f) (Supp. Fig. 7), implying that these estimated false positive rates are generalisable to other samples.

To evaluate the precision of the nanopore-detected modified mRNA sites, we tested the site-level models against sites reported by orthogonal methods on human cell lines. Our results demonstrate that SWARM models trained on all-context sequences performed better when restricted to major writer motifs, recovering more validated sites with higher precision despite testing fewer sites. Specifically, nanopore-detected m6A sites showed higher overlap with GLORI when restricted to DRACH motifs (Supp. Fig. 8a & 8b). This is relevant as some of the current m6A models for SQK-RNA002 only apply to DRACH motifs. Notably, overlap of non-DRACH sites with GLORI was low for all tools and sequencing kits (Supp. Figs. 9a & 9b). When comparing SWARM with other SQK-RNA002 models restricted to DRACH sites, SWARM achieved a higher agreement with GLORI sites in HeLa (Fig. 2g). SWARM also maintained a high overlap with GLORI sites for SQK-RNA004 on DRACH sites, with similar accuracy to dorado (Supp. Fig. 9c). Importantly, at 90% precision, SWARM recalled more GLORI sites than any other available method for SQK-RNA002 or SQK-RNA004, demonstrating top performance of m6A on DRACH sites across both chemistries (Fig. 2h).

Similarly to m6A, we observed that SWARM-detected Ψ sites had higher overlap with bisulfite methods ^30,32^ when restricted to sites with motifs for TRUB1 (GUΨCNA) or PUS7 (UNΨAR, N=A,C,G,U) (Supp. Figs. 8c & 8d), the main Ψ writers modifying mRNA ^35,36^. SWARM also surpassed other nanopore methods in the accuracy of Ψ detection when comparing predicted sites with orthogonal methods BID-seq and PRAISE in HEK293T (Fig. 2i), for both TRUB1 and PUS7 sequence contexts as well as in all sequence contexts (Supp. Figs. 9d-f). Furthermore, SWARM exhibited significantly higher agreement than competing approaches across both chemistries when evaluating the top 100 most confident nanopore predictions with TRUB1 and PUS7 sites detected by bisulfite methods (Fig. 2j), demonstrating best-in-class performance for precise Ψ site detection.

In contrast to m6A and Ψ, all available methods for nanopore m5C site detection showed low overlap with bisulfite sites ^26^ in all sequence contexts. Despite an increase in the overlap when restricted to sites for NSUN6, a major m5C synthase with a well-defined motif (CUCCA) ^37^, m5C site detection with nanopore remains lagging behind m6A and Ψ detection in terms of orthogonal support ^6^ (Supp. Figs. 8e & 8f) (Supp. Figs. 9g-j).

### Limited co-occurrence of m6A and Ψ observed across mammalian tissues

Leveraging SWARM’s specificity and ability to detect modified sites with high precision, we mapped the m6A and Ψ landscapes across six tissues (Cerebellum, Frontal Cortex, Hippocampus, Liver, Testis, Skeletal Muscle) from five mammals (Human, Mouse, Rat, Dog, Cow) ^7^. Given the demonstrated higher precision when restricting to well-established motifs, we focused the analysis on sites potentially associated with METTL3 (DRACH) for m6A and with TRUB1 (GUΨCNANNC) and PUS7 (UVΨAR, V=A,C,G) for Ψ. To select modification sites, we used a 10% stoichiometry cutoff and site-level probability cutoffs that yielded FPR < 0.1% in tests against unmodified cellular mRNA IVTs in those motifs (Supp. Table 3). We detected a total of 376,674 m6A sites (Supp. Table 4) across 79,284 transcripts, and 3,682 Ψ sites (Supp. Table 5) across 3,468 transcripts (Fig. 3a). The consensus sequence of detected sites (Fig. 3b) recapitulated previously established motifs for TRUB1 ^36^, PUS7 ^25^, and METTL3 ^12^ beyond restricted nucleotides, providing confidence for the detected sites.

**Figure 3.**
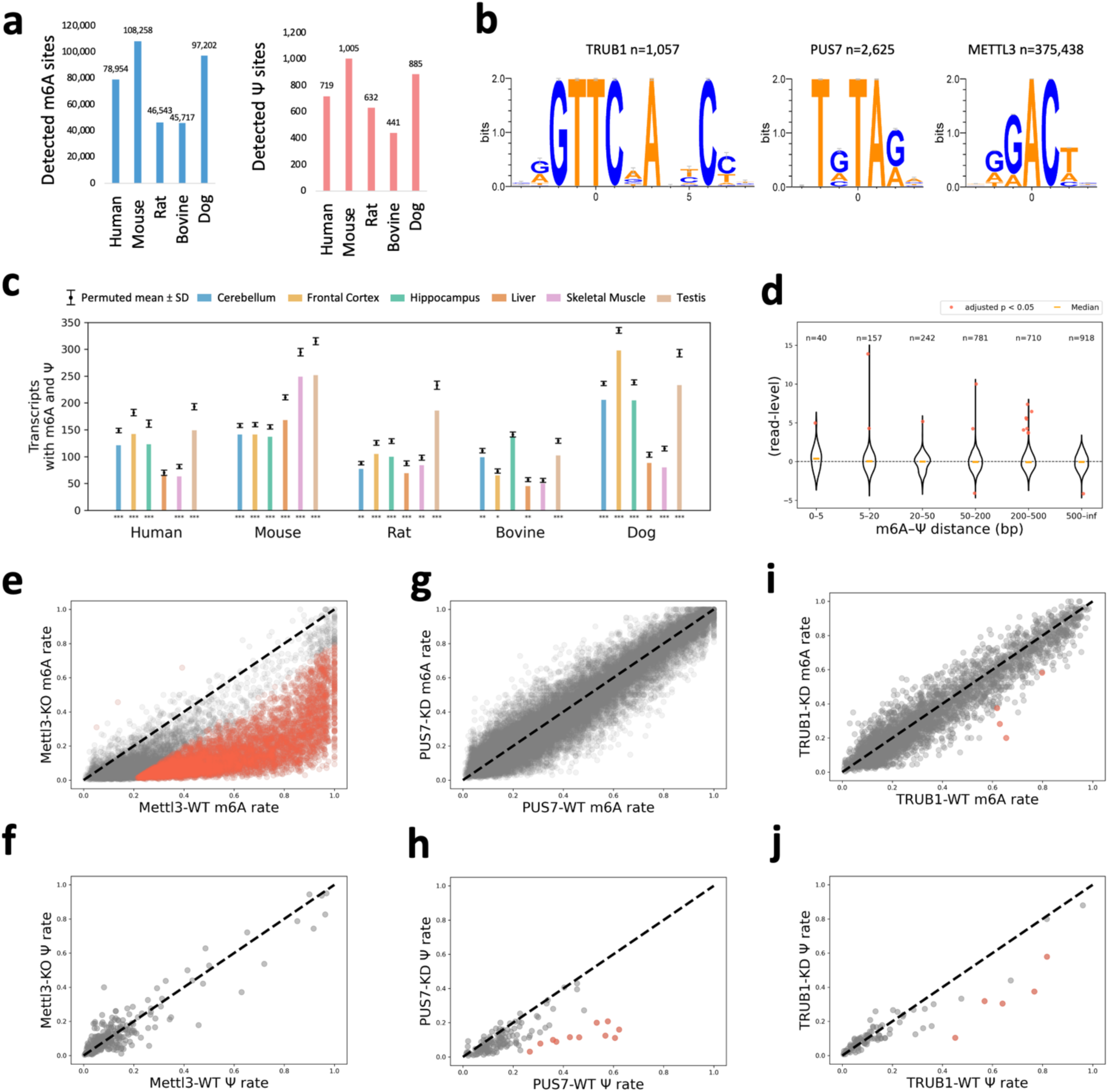
m6A and Ψ co-occurrence in mammalian tissues. **(a)** Number of m6A sites (left panel) and Ψ sites (right panel) detected in at least one tissue of each species. **(b)** Logo plots of detected sites with TRUB1, PUS7, and METTL3 substrate motifs (left to right). **(c)** Number of transcripts with at least one METTL3 candidate m6A site and one PUS7 or TRUB1 candidate Ψ site for each species and tissue (x-axis). On top of each bar, we show the expected number of cases (mean and standard deviation) from permuted modification states across all the tested METTL3 and TRUB1/PUS7 candidate sites in transcripts. Significant p-values are indicated: * = 0.05; ** = 0.005; *** = 0.001. **(d)** For pairs of m6A and Ψ sites in the same transcript at a distance range (x-axis), we plot the z-score distribution obtained by comparing the observed read-level co-occurrence, calculated as the proportion of reads where both modifications are present, with the expected co-occurrence calculated by randomising the read probabilities. **(e-j)** m6A (e,g,i) and Ψ (f,h,j) stoichiometry in KO/KD (y-axes) backgrounds of: HEK293T Mettl3 KO (e-f), HeLa PUS7 KD (g-h), and HEK293T TRUB1 KD (i-j) against the same cell-line control (x-axis). M6A sites restricted to DRACH sites reported by GLORI ^12^, Ψ sites restricted to ones reported by BACS ^25^ with motifs for PUS7 (UNΨAR) and TRUB1 (GUΨ) (f), PUS7 (h), and TRUB1 (j). Sites significant after Benjamini-Hochberg correction (p-value < 0.05) and stoichiometry change > 0.2 are coloured red. Significance was tested using a generalised linear model comparing stoichiometry with observed coverage between the two conditions.

Furthermore, 2,563 unique transcripts across all species harboured both Ψ and m6A sites, comprising on average 1.24% of transcripts per condition. We assessed whether the observed m6A-Ψ co-occurrence within transcripts is different from expected by chance. Considering transcripts with both METTL3 and TRUB1 or PUS7 candidate sites tested by SWARM, we compared the observed co-occurrence with the mean and standard deviation of the overlaps obtained after permuting all sites tested 1000 times. Our analysis indicates that the m6A-Ψ co-occurrence within transcripts is mildly lower than expected, with an average 16% reduction across all tissues and species. The resulting z-scores were consistently negative in all samples, with a statistically significant (p < 0.05) decrease in modification co-occurrence in 27 out of the 30 samples tested (Fig. 3c) (Supp. Fig. 10a).

### No evidence of direct mutual exclusion between m6A and Ψ in predominant writer contexts

Following up on the observed reduction of transcript-level site co-occurrence, we investigated the m6A-Ψ co-occurrence patterns in individual reads. Specifically, we selected m6A-Ψ site pairs based on their closest distance in transcripts where both modifications were detected and compared the observed co-occurrence with the mean and standard deviation obtained after performing 1000 permutations of the same reads covering each pair of transcript sites. Only 16 out of the 2,848 site pairs tested showed a statistically significant difference from expected co-occurrence (adjusted z-score test p < 0.05) (Fig. 3d). In addition to the high proportion of non-significant sites, the distribution of z-scores computed among these sites did not diverge from normality (Supp. Fig. 10b), with a median close to zero (−0.04), supporting the notion that m6A and Ψ occur in individual molecules independently of each other. Interestingly, a similar z-score distribution was observed regardless of the distance between m6A and Ψ sites, with molecule-level co-occurrence appearing random even when the m6A and Ψ sites were close to each other (Fig. 3d).

To further assess the potential association between m6A and Ψ, we analysed the possible differences in m6A and Ψ stoichiometries in METTL3-knockout ^38^, PUS7-knockdown ^39^, and TRUB1-knockdown ^14^ conditions and corresponding controls. Focusing only on sites previously reported by orthogonal methods ^12,25^, we observed a dramatic depletion of m6A in METTL3-knockout samples compared to controls, but no detectable significant differences in Ψ between the same samples (Figs. 3e & 3f). In turn, PUS7 knockdown significantly reduced Ψ stoichiometry, with no significant m6A changes across DRACH sites (Figs. 3g & 3h). Similarly, TRUB1 knockdown induced reduction of Ψ, while only 0.11% of tested sites showed a significant m6A change (Figs. 3i & 3j), indicating a lack of widespread TRUB1 impact on m6A.

### METTL3, TRUB1, and PUS7 candidate sites show distinct distribution patterns along mRNA

We next assessed the distribution of m6A and Ψ sites along transcripts. We annotated the positions of candidate modification sites within UTR (untranslated regions) and CDS (coding sequence) regions using nanopore data from six tissues in five different mammals ^7^. Our detected sites recapitulate the already described m6A distribution, with an enrichment of m6A in the 3’UTR after the stop codon ^12^ (Supp. Figs. 11a-b). The Ψ distribution also peaked in 3’ UTRs, with human PUS7 sites showing strong bias than TRUB1 sites (Supp. Fig. 11c). This recapitulates the trend observed with BACS in HeLa ^25^ (Supp. Figs. 11d-e). Interestingly, detected Ψ sites followed the distribution of sites tested with PUS7 and TRUB1 motifs (Supp. Fig. 11e), suggesting that Ψ deposition may be driven by their cognate motifs. Focusing on the CDS region, m6A sites were enriched in the 2^nd^ codon position compared to the tested sites (Supp. Fig. 11e). Similarly, Ψ sites were found preferentially in the 2^nd^ codon position, with PUS7 sites being significantly enriched compared to the tested sites, similar to the trend observed with BACS in HeLa sites ^25^ (Supp. Fig. 11e).

Furthermore, our high-precision mapping revealed a distinct spatial organisation of Ψ, confirming its localisation peak near exon-exon junctions ^35^. This pattern, observed for Ψ sites with TRUB1 (Fig. 4a) and PUS7 (Supp. Fig. 12a) motifs, contrasts with the well-documented depletion of m6A marks across exon-exon boundaries ^16,40–43^ (Supp. Fig. 12b). In fact, Ψ localisation is largely driven by an overrepresentation of the respective PUS7 and TRUB1 recognition motifs near splice sites (Supp. Fig. 12a). When normalising for this underlying positional bias, the Ψ modification rate (i.e. detected over tested) remains relatively constant across exon-exon boundaries (Supp. Fig. 12c).

**Figure 4.**
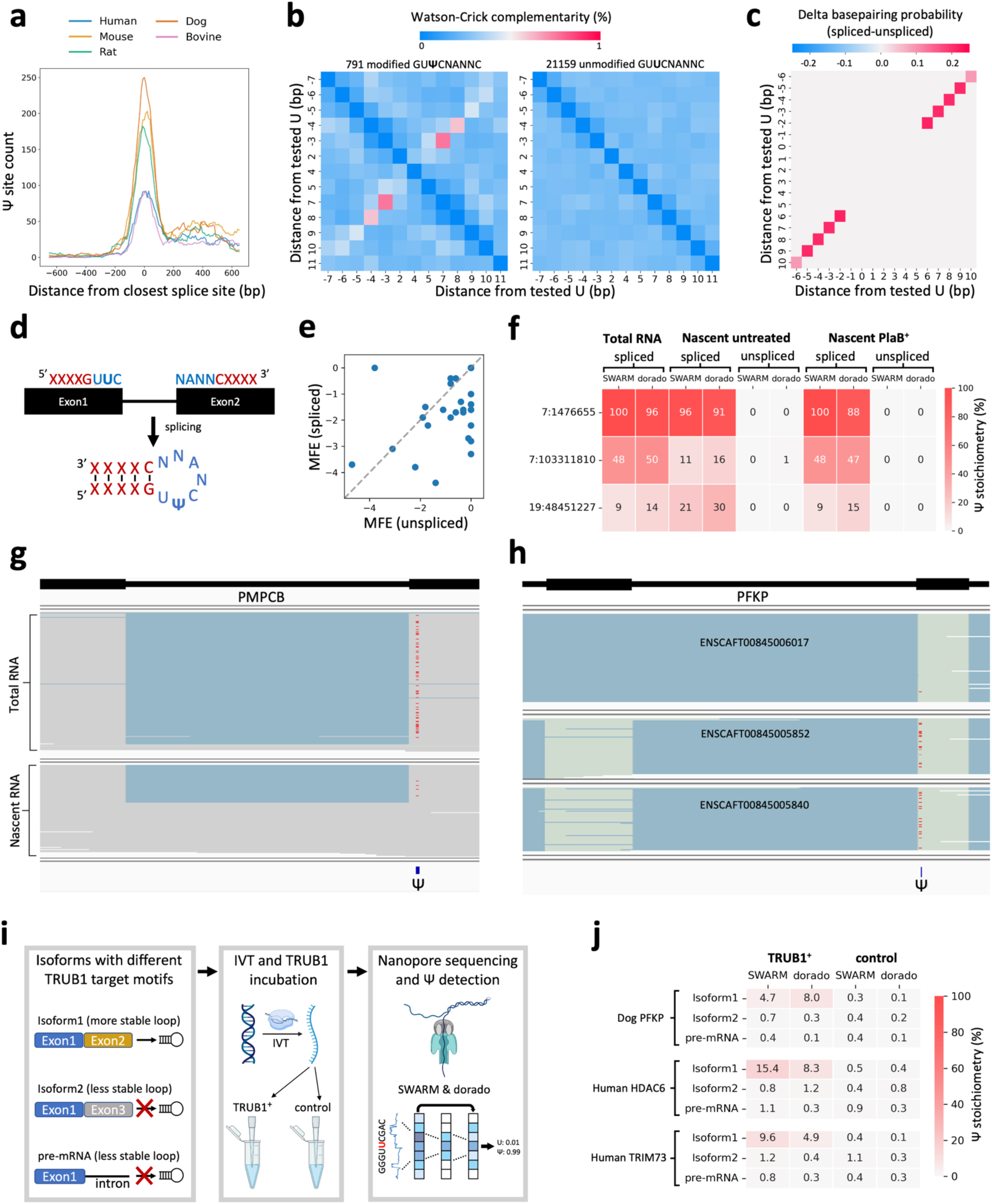
Splicing-shaped TRUB1 pseudouridine sites. **(a)** Counts of Ψ sites detected in each species in 100bp sliding windows centred on the closest exon-exon junction. **(b)** Base-complementary matrix across 791 Ψ sites (left) and 21,159 tested unmodified sites (right) with TRUB1 consensus motifs in 5 mammals and 6 tissues. Unique genomic sites detected in at least one transcript in at least one tissue were analysed in each species. Complementarity was calculated as in Safra et al. ^36^, using only Watson-Crick base-pairing. **(c)** Delta of base-pairing probability values for all positions around the 42 TRUB1-candidate Ψ sites identified in mammals with the substrate structure crossing the exon-exon boundaries, comparing the spliced and the pre-mRNA (i.e., including the adjacent intron) sequence. Unlike in (b), non-Watson-Crick base-pairs were also considered. **(d)** Schematic showing TRUB1 substrate structure formation after splicing. **(e)** Scatter plot showing the minimum free energies (MFE) of the candidate TRUB1 substrate structure in the spliced (y-axis) and pre-mRNA (x-axis) for the 42 mammalian sites where the TRUB1 substrate structure motif crossed the exon-exon boundary. Paired t-test was used to measure the difference in MFE between the pairs of spliced and unspliced sites: n = 42, df = 41, t = −3.107, p = 0.00343, mean difference = −0.6048, 95% confidence interval = [−0.9979, −0.2117], Cohen’s d = −0.4794 **(f)** Ψ stoichiometries in three HeLa sites with the TRUB1 substrate structure interrupted by an exon-exon boundary, measured by SWARM and dorado in three RNA fractions: reads from total RNA sequencing, and spliced or unspliced reads from nascent RNA sequencing. **(g)** IGV snapshot of the human PMPCB gene showing reads from total RNA (upper track) and nascent RNA fractions (lower track) in HeLa. Gray blocks represent positive-strand reference matches, and blue lines represent introns. Below, the position of the Ψ site (7:103311810) is indicated. Read-level Ψ SWARM predictions with probability > 0.9 are shown in red. **(h)** IGV snapshot of the dog PFKP gene in testis tissue samples with three isoforms present in each alignment track. Green blocks represent negative-strand reference matches, and blue lines represent introns. Below, the position of the Ψ site (2:32615598) is indicated. Read-level Ψ SWARM predictions with probability P>0.9 are shown in red. **(i)** Experimental design for the validation of TRUB1-dependent splicing-shaped Ψ sites. **(j)** Ψ stoichiometry measured by SWARM and dorado in TRUB1^+^ and control IVT samples across different splicing isoform constructs of the same gene.

### Splicing-shaped pseudouridylation

The proximity of the Ψ sites to the splice-sites raises the question of whether the substrate sequence for pseudouridylation may be affected by the formation of splicing boundaries, especially for TRUB1, which in mRNA shows affinity for an extended motif that includes a stem-loop similar to the tRNA TRUB1 substrate ^36^. Across the six tissues from five mammals analysed, we detected a total of 791 unique candidate TRUB1 Ψ sites, i.e. Ψ sites predicted by SWARM on GUΨCNANNC sequence contexts. These sites showed an extended motif compared with control sites (tested GUUCNANNC sites but negative for Ψ), suggesting the expected stem-loop structure ^36^ (Fig. 3b) (Supp. Fig. 13). Confirming this, the 791 TRUB1-motif sites showed sequence complementarity between the expected positions, which was absent in the control unmodified sites (Fig. 4b) (Supp. Fig. 14). From these 791 sites, 42 (5.3%) were located close enough to the exon-exon boundary that the expected structure would inevitably cross it. Specifically, the location of the boundary relative to the Ψ site ranged from 5 nucleotides upstream (i.e., Ψ occurs downstream of the boundary) to 9 nucleotides downstream (i.e., Ψ occurs upstream of the boundary). These 42 sites showed higher average base pairing probability in the stem of the TRUB1 substrate structure when the exon-exon sequence was considered (i.e., spliced mRNA) compared to when the exon and adjacent intron sequences were considered (i.e., pre-mRNA) (Fig. 4c), suggesting that TRUB1 Ψ sites nearby exon-exon boundaries preferentially occur after splicing (Fig. 4d). Furthermore, we also detected a significantly lower minimum free energy (MFE) for the TRUB1-substrate structures in the spliced transcripts compared to the pre-mRNA ones (paired t-test p-value = 0.00343) (Fig. 4e). From all the 42 cases detected, 20 showed a more stable structure in the spliced transcript, implying that these TRUB1 sites may only be modified after splicing. For the rest, 15 showed no difference in stability, and 7 showed a more stable structure in the unspliced transcript, suggesting that some TRUB1 substrates may also form before splicing.

To validate the potential link of TRUB1 pseudouridylation with splicing, we used direct RNA sequencing of pre-mRNA from HeLa cells ^44^. From the six Ψ sites detected in human tissues (out of the 42 detected across five mammals), three of them had read coverage in HeLa pre-mRNA. In all three cases, Ψ was detected by SWARM in the spliced reads, but was not observed above detectable levels in the pre-mRNA reads that included the adjacent intron (Figs. 4f & 4g) (Supp. Fig. 15), confirming the expected modification status based on differences in TRUB1 structure stability. As a control, we measured Ψ stoichiometry at sites harbouring TRUB1 motifs fully contained within a single exon. The stoichiometry at those sites was generally lower in pre-mRNA reads compared to total (spliced) RNA reads (Supp. Fig. 16), raising the question of whether modification depletion imight be due to timing. To assess this, we analysed pre-mRNA reads from samples treated for 30 minutes with Pladienolide B (PlaB) to inhibit splicing ^44^. We detected an increase in the modification levels in unspliced reads in most within-exon sites in the PlaB-treated samples compared with the untreated samples (Supp. Fig. 16), indicating that the internal TRUB1 sites had sufficient time to be modified. In contrast, for the three splice-interrupted Ψ sites for which we had sufficient coverage, the modification remained absent in unspliced reads even after PlaB treatment (Fig. 4f). This confirms that TRUB1 sites disrupted by splice-site boundaries can be modified only after exon-exon ligation has occurred.

We next considered whether splicing-shaped pseudouridylation could occur in the context of alternative splicing. To control for potential differences due to tissue context, we focused on alternative splicing events in the same tissue. Out of the 42 splicing-shaped candidate TRUB1 sites detected across mammalian tissues, 20 were located near alternative splicing events calculated from the reference annotation with SUPPA ^45^. However, only one of these sites was captured in multiple isoforms expressed in the same tissue, a Ψ site in the *PFKP* gene in dog testis near an exon skipping event, with Ψ detected in reads including the alternative exon, and Ψ depleted in reads skipping the exon (Fig. 4h). Confirming the splicing-shaped nature of this site, we confirmed a higher stability of the TRUB1 substrate structure in the reads including the alternative exon compared with the reads skipping the alternative exon (Fig. 4h) (Supp. Table 6).

Since read coverage of alternative isoforms may limit the ability to observe splicing-shaped pseudouridylation, we searched for additional sites that could be predicted by scanning the reference annotation. We identified a total of 1412 hypothetical TRUB1 Ψ sites in the human transcriptome, i.e. GU**U**CNANNC was found in a transcript isoform with base-pair complementarity to form the TRUB1 substrate, which is a strong predictor of TRUB1 site modification ^36^. Of those, 69 (4.8%) (including the 6 observed in the human tissues with SWARM) had a TRUB1 structural motif intercepted by a splice site. Moreover, 20 of these candidate Ψ sites showed potential for regulation by alternative splicing (Supp. Table 6), occurring within annotated alternative isoforms that showed sequence differences in the TRUB1 binding sequence that would disrupt the substrate structure.

To further validate splicing-shaped pseudouridylation, we generated IVT constructs corresponding to mRNA isoforms with different TRUB1 substrate motifs and their pre-mRNA sequence for five different genes, using valine tRNA as a positive control (Fig. 4i) (Supp. Table 7). After incubating the IVT RNAs with TRUB1 (Methods), Ψ was observed with high specificity for the expected loci in each gene construct (Supp. Fig. 17), and all five tested genes showed variation in Ψ levels across the different isoforms, confirming the splicing-shaped nature of TRUB1 pseudouridylation in mRNA. Specifically, we confirmed that the site observed in dog *PFKP* gene described above has detectable Ψ levels at the target site only in the construct for the spliced isoform with a more stable TRUB1-substrate structure (Fig. 4j) (Supp. Table 7). Additionally, we tested two human sites that, while not detected in the nanopore data, were predicted from sequence and structural analysis. In both these cases, only the isoform with a more stable structure was modified (Fig. 4j) (Supp. Table 7). We also tested a site in the cow gene *IFT22,* for which we observed the opposite trend in the candidate TRUB1-substrate in the cow annotation, with higher structural stability in the pre-mRNA. For this case, we validated the highest Ψ levels in the construct corresponding to the pre-mRNA (Supp. Table 7), confirming that, as suggested by our analyses above (Fig. 4e), some TRUB1-dependent sites may be modified prior to splicing. Finally, we tested a site in the human *PMPCB* gene, assessing the canonical isoform ENST00000249269, where Ψ was observed in previous experiments, the exon-skipping isoform ENST00000706440, which had no coverage in cellular data but had a more stable TRUB1 substrate structure, and the intron-retaining isoform ENST00000706438 (Supp. Table 7). As expected by the structural analysis, the exon-skipping isoform ENST00000706440 showed high Ψ levels after TRUB1 incubation compared to the canonical isoform. Surprisingly, the *in vitro* experiment for ENST00000249269 yielded opposite results to the cellular measurements showing higher modification levels in the intron-retaining construct compared to the spliced canonical isoform construct. When we considered the RNA structures of the IVT constructs, we observed that the known TRUB1 site was occluded in the construct corresponding to the canonical isoform but was more accessible and closer to the TRUB1-cognate structure in the intron-retaining construct (Supp. Fig. 18). This shift in modification patterns in the absence of the nuclear environment suggests that the presence of other factors, such as spliceosomal complexes and splicing regulators, may shape the TRUB1-dependent Ψ landscape.

## Discussion

SWARM provides a significant methodological advance in epitranscriptomics by establishing a robust, high-resolution workflow for the simultaneous detection of m6A, Ψ, and m5C from individual nanopore RNA sequencing reads. A key innovation in SWARM is its training strategy, which integrates synthetic RNAs with target and non-target modifications to improve modification specificity, as well as signals from orthogonally validated cellular sites to achieve high-precision detection of modified sites transcriptome-wide. Our training strategy effectively mitigates the confounding influence of neighbouring modifications on nanopore signals and results in a drastic reduction in false positives, a significant limitation of existing tools. This capability is a fundamental leap beyond existing tools, as demonstrated by SWARM’s superior performance in rigorous benchmarking against diverse independent datasets. We note that modification crosstalk can also occur when different modification types are densely localised within the same RNA. This aspect of crosstalk was not tested due to the lack of appropriate training and testing data, but it is an interesting avenue for future studies.

A clear outcome of our comprehensive benchmarking is the exposure of a major methodological gap: the poor concordance between nanopore sequencing and orthogonal methods for m5C detection. This likely reflects both the biological nature of m5C, such as low stoichiometries and highly context-dependent deposition, as well as technical inconsistencies among assays. While SWARM does not resolve these challenges, it provides a reproducible, high-standard benchmark upon which future methodological developments can build. This could also serve as a catalyst for the community to develop validated controls and orthogonal standards for m5C, a necessary step for reliable epitranscriptome discovery ^46^. The development of comprehensive standardised benchmarking datasets could help improve the detection of other modifications using nanopore sequencing and other technologies ^47^, especially rare ones or those that appear only under very specific conditions ^6^.

The modular design of SWARM is a key strength, providing a framework that can be extended to incorporate additional RNA modifications as additional high-quality training and testing datasets for cellular RNA become available. Moreover, unlike most other detection tools, SWARM is compatible with both SQK-RNA002 and SQK-RNA004 chemistries, ensuring broad applicability to new and legacy datasets. We have illustrated this capability with our application of SWARM to diverse mammalian transcriptomes, a rich nanopore-based resource generated with the SQK-RNA002 chemistry ^7^. The unique SWARM’s accuracy on SQK-RNA002 data for m6A and Ψ enabled us to challenge existing paradigms about modification coordination and to reveal new principles of pseudouridylation.

A significant finding of our study is the absence of direct coordination between m6A and Ψ modifications in the predominant writer contexts. Despite their independent enrichments in mRNA and potential roles in regulating mRNA metabolism, our analyses of the transcript and read-level co-occurrence and perturbation experiments consistently showed no evidence of a strong, widespread positive or negative association. However, the slight but significant underrepresentation of transcripts with both modifications suggests these marks may be subject to selective pressures favouring their mutual exclusivity, perhaps to avoid conflicting regulatory outcomes. While these findings challenge the emerging hypothesis of a widespread coordinated epitranscriptomic code, they do not rule out the possibility that specific combinations of modifications might be co-regulated or define more nuanced functional outcomes in specific mRNAs. Furthermore, modification sites corresponding to the less predominant m6A and Ψ writers, which were not tested in this study, require further analysis and remain subject to unknown co-occurrence patterns. Tools like SWARM can help dissect these complex relationships.

Our analyses also provided compelling evidence for the splicing-shaped TRUB1-mediated pseudouridylation. We identified Ψ sites near exon-intron boundaries whose predicted local RNA structures become significantly more stable in the mature, spliced mRNA than the pre-mRNA context. We validated this splicing-shaped pseudouridylation using nascent RNA nanopore sequencing as well as with *in vitro* constructs encoding different exon-exon and exon-intron isoform variants, where pseudouridylation was evident in the spliced isoforms but absent or depleted in the unspliced transcripts or alternatively spliced variants with less stable TRUB1-cognate RNA structure. Interestingly, although less frequent, we also identified and validated cases where Ψ is deposited at a higher rate in the pre-mRNA context when there was a stable TRUB1-cognate RNA structure already present. Due to read coverage limitations and rare nature of Ψ, we were able to show experimental evidence for the splicing-shaped pseudouridylation for seven unique pseudouridine sites, and we confirmed TRUB1 dependency for five of these sites using *in vitro* experiments. Our results collectively support a novel Ψ deposition mode in which splicing remodels the local RNA structure near splice sites, tuning TRUB1 substrate affinity and facilitating context-dependent Ψ deposition. Our estimates from sequence and structural motifs and transcript annotations indicate that 20 human Ψ candidate sites show potential for modification tuning mediated by alternative splicing. However, the molecular mechanisms underlying the splicing-shaped deposition of Ψ sites and their functional relevance warrant further investigation.

Recent studies suggest that biologically robust nanopore-based detection of RNA modifications is limited to m6A ^23^. The unique nature of m6A, being modified predominantly by a single complex and being frequent in mRNA with overall high stoichiometry, undeniably elevates the capacity of nanopore methods to discover m6A biology with a low amount of noise contamination. In contrast, Ψ in human can be placed by up to 13 different synthases with diverse affinities, and is generally present at low stoichiometry in mRNA, making Ψ detection considerably more prone to noisy predictions. In our study, we demonstrated that a training strategy focused on improving detection precision is highly potent for biologically robust Ψ detection, evidenced by our discovery and experimental validation of a newly identified splicing-shaped mode of TRUB1-dependent pseudouridylation. Our high-precision training paradigm paves the way to achieve biologically robust nanopore analysis of low-frequency RNA modifications.

Our systematic benchmarks indicate that no single RNA modification model is uniformly optimal for all applications. The benchmarks for different SWARM model hyperparameters support this: certain models rank highest for modification-type detection in IVT tests but rank lower for mRNA site detection, and vice versa. Our results indicate that SWARM strikes a good balance, consistently ranking as the highest or among the top tools on the tested tasks for m6A and Ψ detection. For instance, dorado and SingleMod produced a higher m6A stoichiometry correlation with GLORI, yet SWARM had better modification separation and higher precision in the detection of mRNA sites. This behaviour aligns with the training strategy, as SingleMod training was optimised for correlation with GLORI stoichiometry, while SWARM model hyperparameters were selected to optimise read- and site-level specificity.

Furthermore, we strongly advocate that the academic community develop open-source nanopore RNA modification models, with transparent code and training data, to ensure reproducibility and enable rigorous evaluation of model generalisation. This is a major limitation of dorado models, which are trained with undisclosed data, meaning that it is not possible to tell if models have been trained or optimised using signals from the same publicly available data used for model benchmarks, leading to a potential data leakage problem. Consequently, the only way to benchmark dorado models with minimal risk of leakage would be to use in-house data containing unusual modification contexts. However, such datasets are difficult to obtain, and these benchmarks might not be broadly relevant for the research community, which studies the cellular contexts already well represented in public datasets.

In summary, SWARM equips the community with a validated, high-precision workflow to decode the complex landscape of RNA modifications from individual nanopore reads. Its crosstalk-aware design, compatibility with multiple chemistries, and control-free operation make it uniquely practical and robust for mining existing datasets and designing new experiments. As an open-source tool with standardised benchmarks, SWARM provides an indispensable resource for advancing our understanding of the epitranscriptome in health and disease.

## Methods

### Preparation of synthetic RNAs and nanopore direct RNA sequencing

Reference *in vitro* transcribed RNAs (IVTs) i1-i4 and m1-m4 ^16^ were obtained by run-off in vitro transcription from plasmids encoding bacteriophage T7 RNA polymerase promoter followed by a unique artificial sequence covering the entire length of the desired RNAs (plasmids were ordered in 50 μg aliquots from GeneScript Biotech). Eight pUC57-mini-X (X = i1-i4, m1-m4) DNA templates were generated by plasmid linearization with different restriction endonucleases. The plasmids and the restriction digestion protocol are identical to those used previously ^16^. Briefly, the i1-i4 sequences contained random 5-mers, whereas m1-m4 sequences were built from four non-overlapping fragments of the mouse canonical pre-rRNA (∼13kb long) (Supp. Table 1).

The *in vitro* transcription reaction was conducted as previously described ^16^. For each transcription reaction, 1 μg of each IVT was mixed with nucleotide buffer mix (NTP) to yield a final concentration of 10 mM of each nucleotide, 1 unit of RNasin Plus (Promega), 5 mM of dithiothreitol (DTT) and 2 units of T7 RNA polymerase mix with the total reaction volume scaled to 40 μl. The transcription reaction mixture was incubated at 37°C for 2.5 hours, after which the sample was diluted to 100 μl using deionised water. The diluted sample was then mixed with 1× DNase I buffer (Thermo Fisher Scientific) and 2 units of DNase I (Thermo Fisher Scientific) and was incubated for 15 minutes at 37°C. Following the 2.5-hour reaction time, 2.5× volumes of AMPure XP SPRI bead suspension (Beckman Coulter) was added to this mixture, and binding was conducted for 10 minutes at 25°C with periodic mixing. As recommended by the bead supplier, the binding step was followed by two sequential washing steps each with 1 ml of 80% v/v ethanol. The RNA sample was eluted in 30 μl of deionised MQ water. The concentration of the RNA samples was assessed initially with nanodrop spectrophotometer, and its quality was additionally assessed with Qubit Broad Range (BR) assay using Qubit spectrophotometer (Thermo Fisher Scientific).

For the *in vitro* polyadenylation reaction, RNA was preheated at 65°C for 2 min prior to polyadenylated by E. coli poly(A) polymerase (New England Biolabs). Approximately 5µg of RNA, 6µl of EPAP, 1mM ATP, 2µl of RNasIn, and 2µl of T7 RNA polymerase (all components purchased from New England Biolabs), were incubated for 30 min at 37°C water bath. The poly A-tail containing RNA was purified with 1 ml freshly prepared 80% v/v ethanol and AMPure XP beads (Beckman Coulter). The polyadenylated RNA was subsequently used for the library preparation and sequencing using ONT direct RNA sequencing protocol as described below.

The sequencing library was prepared as per the manufacturer’s recommendation: 1000ng of RNA was used for 2X library preparation for the IVT construct. The RNA Control Standard (RCS) was eliminated from the ligation reactions. Superscript IV and RNAsin were additionally introduced for ligation incubation and reverse transcription. The sample was purified with 2:1 ratio of sample to AMPure XP beads and wash with 80% freshly prepared ethanol. The sample went through a second incubation with NEBNext quick ligation buffer and T4 DNA ligase (New England Biolabs) for 15 minutes at room temperature. The RNA sample was further cleaned up with AMPure XP beads and wash buffer provided by manufacturer (ONT). Then, the reverse transcribed and adapted RNA was eluted using 20-40 μl elution buffer (ELB). The direct RNA sequencing was conducted using both pore chemistries, SQK-RNA002 and SQK-RNA004. For SQK-RNA004, the full protocol was identical to the SQK-RNA002 except for minor adjustments for the reverse transcriptase reagents and incubation time: ProtoScript II was used for the reverse transcription reaction: 25°C for 5 minutes, then 42°C for 60 minutes, 65° C for 20 min, and bringing the sample temperature to 4° C.

For library loading, the flow cell was first inserted under the MinION clip to ensure proper thermal and electrical contact. Quality control of the flow cell, to assess the number of pores, was performed, and the priming port was opened to remove any air bubbles. A priming mix, comprising of the RNA Flush Tether (RFT) and Flush Cell Flush (FCF) (Activated Flush Buffer), was prepared and loaded into the flow cell as per manufacturer’s instructions. Following this, the loading library was prepared by mixing the ELB eluted RNA with Sequencing Buffer (SB) and Library Solution (LiS). Upon mixing, the library was gently loaded into the flow cell via the SpotON sample port in a dropwise manner. After loading both the activated flush buffer and the library into the FLO-PRO004RA flow cell, the SpotON sample port cover was replaced, the priming port was closed and MinION was secured in place, following the recommended guidelines. The DRS sequencing runs were conducted at room temperature (∼25 °C) up to a maximum duration of 72 hours with wash and reload, where the second half of the library was loaded for an additional 24 hours.

### Synthesis, amplification, and nanopore direct RNA sequencing of cDNA libraries

15 ng of Human Brain, Hippocampus Marathon®-Ready (Catalogue number *639319*) and Mouse 7-day (D7) Embryo Marathon®-Ready (Catalogue number *639407*) were purchased from Clontech (Takara Bio, USA). All procedures relating to the reverse transcription of poly(A) enriched RNA to the synthesis of Marathon-Ready cDNA for these samples can be found in Marathon®-Ready instruction manual by the manufacturer. cDNA amplification of the samples was conducted to increase the yield of the starting amount of cDNA from 15 ng to at least 1 mg, sufficient for RNA production and to generate very high-yield libraries with as many diverse sequence contexts for SWARM training models. The cDNA amplification procedure to amplify human hippocampus and mouse embryo D7 cDNA samples was adapted from the Marathon® end-to-end PCR procedure (Takara Bio, USA).

To set up the amplification, AP1 primer was provided from the Marathon® kit to be used as the negative control, as recommended by manufacturer protocol. Since all sequences in the pool of cDNA samples harboured the reverse complement sequence of AP1 adaptor nucleotides at the 5’ end, we designed in-house primers of AP1-F: *CTAATACGACTCACTATAGGGCTCGAGCG* (T_m_ = 67.5°C) as the forward primer and AP1-R: *TTCTAGAATTCAGCGGCCGCTTTTTTTT* (T_m_ = 67.5°C) as the reverse primer, designed around the poly(A) tail of adaptor sequence. These primers were used to amplify all transcripts in the cDNA sample. The primers were designed and ordered from Integrated DNA Technologies (IDT), where 100 mM of master stock solutions of each forward (AP1-F) and reverse primers (AP1-R) as well as the original AP1 primer were made in deionised mQ water.

To prepare the PCR master mix for each cDNA sample, 5 mM of AP1-F and AP1-R were mixed with 1×SuperFi II buffer and Platinum SuperFi II Polymerase (Thermo Fisher Scientific), supplemented with dNTPs added to a final concentration of 200 mM. The total PCR master mix was prepared in a final volume of 50 μl. A parallel negative control for each cDNA sample was set using the same conditions as described above, except that the primers were replaced with the kit-provided AP1 primer.

The PCR master mix for each cDNA sample was incubated in a PCR thermocycler with preset amplification conditions. The steps involved denaturation at 98°C for 30 seconds, followed by annealing step at 60°C for 30 seconds, and the final extension step at 72°C for 4 minutes. The entire protocol is set to 25 cycles. The amplification was confirmed by running the samples on a 2% agarose gel to visually confirm for the presence of bands indicating successful amplification of the cDNA samples. Upon visually confirming the presence of cDNA amplification, the samples were further purified by adding AMPure XP SPRI bead suspension (Beckman Coulter) to the cDNA samples following the same steps as for the IVT samples.

Following amplification, *in vitro* transcription of both cDNA samples was performed using HiScribe T7 High Yield RNA Kit (New England Biolabs) following manufacturer’s instructions and recommendations. The transcription reaction and DNase treatment were conducted as previously described above for the IVT samples. The sample was diluted to 100 μl using deionised water.

*In vitro* polyadenylation of the human hippocampus and mouse embryo D7 RNA samples was performed following the transcription reaction as described above as in IVT samples. A similar RNA extraction process, as described above in the transcription reaction, was repeated. Two independent replicates of each sample were separately subjected to the *in vitro* polyadenylation reaction.

Poly(A) enriched RNA samples from human hippocampus and mouse embryo D7 for direct RNA sequencing (DRS) were prepared using the SQK-RNA004 chemistry, according to ONT protocol, with minor alterations. The same protocol described above was used for the IVT samples. All libraries were prepared in double volumes of the SQK-RNA004 protocol unless otherwise indicated. The only deviation was that the Induro RT reaction was conducted for reverse transcription at 55 °C for 10 minutes, followed by 4 °C hold in the PCR thermocycler. All incubation durations, concentrations, and temperatures were rigorously maintained in accordance with SQK-RNA004 protocol provided by ONT throughout the entire duration of the library preparation protocol. All cDNA libraries, including replicates, were sequenced using a PromethION device and PromethION Flow Cells (FLO-PRO004RA flow cells). Live basecalling was disabled during the run, and the file format for data acquisition was set to POD5 for SQK-RNA004 and FAST5 for SQK-RNA002 for subsequent downstream analysis.

### Targeted tRNA direct RNA sequencing

Total RNA was extracted from 12×10^6^ HeLa cells using standard trizol extraction followed by isopropanol precipitation. Residual DNA was digested using 1 µL of Turbo DNase (Invitrogen, AM2238) in a 40 µL reaction with 2 µL of SUPERase·In™ RNase Inhibitor (Invitrogen, AM2696) for 15 minutes at 37 °C. RNA was re-purified using phenol-chloroform extraction and eluted in 20 µL of nuclease-free water. The purified RNA was run on a 5% w/v polyacrylamide gel containing 7M urea and 0.5x TBE for approximately 1 hour. The region of gel between the xylene cyanol and bromophenol blue bands (tRNA-sized RNAs) was excised and incubated in 0.5M NaCl + 0.01% SDS buffer overnight at 4 degrees in a 1.5 ml total volume. RNA was purified from the eluate and *in vitro* polyadenylated, as described previously ^44^. Approximately 300 ng of polyadenylated tRNA was used for direct RNA sequencing on a PromethION P2 Solo, using SQK-RNA004 library preparation with Induro reverse transcription (NEB, M0681S), as described earlier ^44^.

### Experimental validation of splicing-shaped TRUB1 candidate Ψ sites

DNA templates for *in vitro* transcription were obtained from Integrated DNA Technologies (IDT) as synthetic gBlocks, each containing a T7 RNA polymerase promoter sequence at the 5ʹ end. Barcode sequences and poly(A) tails were introduced at the 3ʹ end of the templates by PCR amplification. Different hexamer barcodes were used for TRUB1-treated (GCATAG) and control (CGATGC) constructs. The reverse primer was designed to anneal to the 3ʹ terminus of the template and to incorporate barcode and poly(A) sequence elements. PCR amplification was carried out using a hot-start Q5 High-Fidelity DNA Polymerase (New England Biolabs). Amplified products were purified using the Wizard® SV Gel and PCR Clean-Up System (Promega) and subsequently used as templates for *in vitro* transcription.

RNA transcripts were generated from the PCR-amplified DNA templates using the T7 RiboMAX™ Express Large Scale RNA Production System (Promega), following the manufacturer’s instructions. Briefly, 1 µg of each DNA template was incubated in a 20 µL reaction containing 10 µL of RiboMAX™ Express T7 2× Buffer and 2 µL of T7 Express Enzyme Mix. Reactions were incubated at 37 °C for 3 h to allow in vitro transcription. Following transcription, RNA products were purified and concentrated using the Monarch® Spin RNA Cleanup Kit (New England Biolabs). RNA concentration was quantified using the Qubit™ RNA Broad Range (BR) Assay Kit (Thermo Fisher Scientific). Purified RNA transcripts for experimental and control conditions (excluding tRNA samples) were pooled separately, with each pool comprising RNAs carrying distinct barcode sequences positioned upstream of the poly(A) tail.

Pseudouridylation assays were performed using four RNA substrates: (i) an experimental RNA pool, (ii) a negative control RNA pool, (iii) a positive control RNA (Val tRNA), and (iv) a Val tRNA mock-treated control. For each condition, 30 pmol of RNA was diluted to a final volume of 40 µL with nuclease-free water. RNA samples were denatured at 75 °C for 2 min and immediately snap-cooled on ice for 5 min. RNA refolding was carried out by the addition of 10 µL of 5× pseudouridylation buffer (final 1× concentration: 100 mM Tris–HCl pH 8.0, 100 mM ammonium acetate, 5 mM MgCl₂, and 0.3 mM EDTA), followed by incubation at 37 °C for 20 min. *In vitro* pseudouridylation reactions were assembled to a final volume of 500 µL by combining 90 µL of 5× pseudouridylation buffer, 10 µL of 100 mM DTT, refolded RNA (46.5 µL), recombinant human TRUB1 (OriGene) to a final concentration of 600 nM, and nuclease-free water. The experimental RNA pool and the Val tRNA positive control were subjected to enzymatic pseudouridylation, whereas the negative control RNA pool and Val tRNA mock control were processed in parallel under identical conditions but without enzyme addition. Reactions were incubated at 30 °C for 45 min. Following incubation, RNA products were purified and concentrated using the Monarch® Spin RNA Cleanup Kit (New England Biolabs) and subsequently used for nanopore sequencing.

### Basecalling and alignment

Reads from the synthetic and cDNA IVTs and the cell lines were basecalled using guppy 6.4.6 configured with the rna_r9.4.1_70bps_hac model for SQK-RNA002, or with dorado version 0.7.2 with the rna004_130bps_sup@v5.0.0 model for SQK-RNA004. Basecalling was accelerated using BLOW5 files ^48^ and buttery-eel ^49^. Basecalled reads were aligned to references using minimap2 ^50^ with parameters: -ax map-ont -k 5 for the reads from the synthetic IVTs, and -k 14 for all other samples. Signals from the synthetic IVT reads were mapped to the sequence templates, whereas all other reads (from cell lines, cDNAs, and mammalian tissue) were mapped to the Ensembl annotation (cDNA references, version 109). The reference used for analysing rRNA reads included the human cDNA reference and GenBank accession numbers U13369 (for 28S rRNA nts 7935-12969), X03205 (for 18S rRNA), U13369 (for 5.8S rRNA nts 6623-6779). tRNA reads were mapped to hg38 high-confidence mature tRNA sequences obtained from GtRNAdb ^51^ (release 22). Event alignment was performed using f5c ^52^ with parameters: f5c eventalign --rna --signal-index --scale-events --samples --print-read-names --sam. with significant speed optimisation and compact storage of aligned signal events in SAM format. ONT 5-mer tables were used for SQK-RNA002 (https://github.com/nanoporetech/kmer_models), while poregen ^53^ 5-mer tables were used for SQK-RNA004 obtained from: (https://raw.githubusercontent.com/hasindu2008/f5c/v1.3/test/rna004-models/rna004.nucleotide.5mer.model).

### Read-level features processing

SWARM parses SAM files with aligned nanopore signal events for every 9-mer sequence context (five consecutive 5-mers) across all provided reads. This context was chosen because the pore models used during event alignment were based on 5-mer sequence segments, and it is accepted that at least 5 bases fit into the nanopore at a given time, meaning that 5 consecutive 5-mer events capture a given base during its pass through the pore. For every base of interest, a 180×7 vector is generated by interpolating scaled signal events for each 5-mer into fixed-length vectors (n The algorithm used by f5C produces outputs identical to nanopolish =36) and using 7 feature channels including: (1) interpolated signal events, (2-5) four channels holding the distances from the expected k-mer signals varying the central 9-mer base (A/C/G/T), (6) dwelling times, (7) per-base q-scores. Signal vector length of 36 for each 5-mer was chosen due to similarity to the median dwelling time, as well as 36*5 being divisible by 9, required for encoding nine q-scores for each base of the 9-mer into the convolution channels. To match the signal feature shapes, dwelling time and q-score features were repeated to add up to the vector shape = 180, meaning that each dwelling time for a 5mer was repeated 36 times, and each q-score from the 9-mer bases was repeated 20 times. Expected event mean signal values were obtained from the same pore models used for event alignment. Signal features are computed using C++ and directly streamed for parallel neural network prediction, circumventing long-term storage of intermediate files.

### Read-level training datasets

Bespoke training datasets were curated for each sequencing kit and target modification. Every training dataset contained up to 500 signals per 9-mer from the all-5-mer synthetic constructs (IVT-i) with IVT sample harbouring target modification for positive data, and an equivalent number of negative signals comprised of 0/20/40/60% combination of equally represented IVT signals from samples with non-target modifications (Ψ, m6A, m5C, ac4C) and remaining percentage of signals from unmodified IVT samples. As 20% non-target modification negatives were selected as optimal on SQK-RNA002 models, SQK-RNA004 models were only trained with the same percentage. While we did not provide a model for ac4C prediction, we generated ac4C IVT data to use as additional non-target modification during training, with the goal of improving modification specificity.

Training dataset for m6A included above mentioned IVT-i signal mixture, supplemented with positive HEK293T mRNA signals (up to 5000 per reference position) from sites with high stoichiometry (>80%) reported by GLORI ^12^, and the same number of negatives originating from the same reference positions in fully unmodified HeLa mRNA IVT ^34^ for SQK-RNA002, and HEK293T mRNA IVT ^54^ + Human Hippocampus IVT (this study) for SQK-RNA004. Dataset for training Ψ models was composed of IVT-i mixtures supplemented with modified signals (up to 5000 per reference position) from rRNA Ψ sites reported by mass-spec ^24^ and BID-seq ^30^ with high stoichiometry (>80%) in an *in vitro* polyadenylated HeLa DRS sample (this study), and negative counterparts in the same number from equivalent positions from unmodified IVT sample synthesised from the mouse rRNA reference (IVT-m) ^16^. In addition to rRNA, the SQK-RNA004 Ψ dataset contained tRNA modified signals (up to 5000 per central 5-mer) from an *in vitro* polyadenylated HeLa sample (this study) at Ψ sites with high stoichiometry (>80%) reported by BACS ^25^, and negative signals within equivalent 9-mers from human hippocampus IVT (this study).

The training dataset for m5C for the SQK-RNA002 chemistry only contained IVT-i signal mixtures, while for the SQK-RNA004 chemistry it additionally included signals from an *in vitro* polyadenylated HeLa sample (this study) (up to 5000 per central 5-mer) at high-stoichiometry sites (>80%) reported by UBS-seq ^26^, and negative signals within equivalent 9-mers from human hippocampus IVT (this study).

Each dataset was split into training/validation/testing using a 60/20/20 ratio, with stratified sampling ensuring the same proportions of 9-mer sequences in each split. In addition, five sub-classes were defined, which comprised different signal contexts and were used to compute sample weights for balancing contributions of each context. Namely, IVT-i-multi, containing multiple target modifications in the same 9-mer, IVT-i-single, with only one target modification in the 9-mer, as well as mRNA, rRNA, and tRNA signals, where applicable.

### Read-level network architectures and training

Considering differences in signal patterns and training data composition specific to each modification, three distinct neural network architectures were trained for each modification, labelled according to the number of parameters (Mini/Mid/Large). Specifically, the *Mini* network was a custom ResNet ^55^ with 5 residual blocks and 96k parameters, the *Mid* network was based on the Jasper architecture ^56^ (128k parameters), and the *Large* network was a customised MobileNet ^57^ with variable filter size (3,7,11) and 6.7m parameters. Each neural network was trained using Tensorflow 2.15.0 with an Adam optimiser for 100 epochs with early stopping and cyclic learning rate. The best epoch was selected based on the minimal validation loss. Each model was trained using either no weights (w0) or balanced weights for all four above-mentioned sub-classes (w1). Models were trained on the NCI gadi high-performance cluster using 4 x GPU Volta chips and 2 x 24-core Intel Xeon Platinum 8268 (Cascade Lake) CPUs. Mini architecture was trained for up to 2 hours, Mid architectures up to 3 hours, while Large architecture required up to 11 hours of training. Inference was computed on individual GPU volta chips with 12 CPUs.

### Site-level model

A site-level model was trained for each read-level model by generating artificial mixtures of modified and unmodified IVT-i signals independent from the training dataset. For generating artificial sites, pools of up to 5000 read-level predictions per reference position were assembled from each of the target modification and unmodified IVT samples. For each reference site, 80 artificial sites were generated with random coverage uniformly sampled between 10-1000 reads, with 50/50 split of positive and negative sites. Positive sites contained a mixture of positive and negative signals with random stoichiometry uniformly sampled between 10% and 100%. Negative sites were generated using primarily unmodified IVT signals at a given reference site, with the additional noise added, including a random sample of 0-4% of predictions from a sample with target modification at the same site, and an identical proportion of random probabilities between 0.2 and 0.8 to represent less confident and noisy read-level predictions. The artificial site mixtures were split 60/20/20 for training/validation/testing.

Input features for all site-level models contained vectors of 100 values representing binned counts of read-level probabilities. All site-level models were trained using the previously described Mini network with the Adam optimiser for up to 100 epochs. Early stopping and a cyclic learning rate schedule were applied, and the best-performing epoch was selected based on the lowest validation loss.

### Model selection

Optimal neural network architecture and training sample weights were chosen for each model based on both modification specificity on synthetic IVT controls and overlap of detected sites with orthogonal methods. Modification accuracy was measured as area under the precision-recall curve (AUPRC) computed on synthetic IVT-i signals from the test split with balanced weights for each non-target modification context. Overlap with orthogonal methods was measured as AUPRC using site-level probabilities evaluated against polyA+ site labels from orthogonal methods: GLORI ^12^ sites for m6A models (excluding training sites) in HEK293T, BACS ^25^ sites for Ψ models in HeLa, and UBS ^26^ sites for m5C models in HeLa. Since most models consistently displayed high AUPRC in IVT controls, models with the highest site overlap with orthogonal methods were selected, with the exception being the m6A model for SQK-RNA004, which was selected according to the highest AUPRC in IVTs, given the small differences in orthogonal overlap and the observed variability in performance on IVT controls.

### Modification specificity benchmarking

Available nanopore modification models were applied to IVT-m samples (IVT-m1, IVT-m2, IVT-m3, IVT-m4), independent from the training data constructs and generated from Mouse rDNA templates containing Ψ, m6A, m5C, and no modifications. Due to coverage constraints, models for SQK-RNA002 were benchmarked on IVT-m1 and IVT-m2 constructs only, while models for SQK-RNA004 were benchmarked using the IVT-m4 construct. Up to 500 read-level predictions per reference position were sampled, and all models were benchmarked using the signals with target modification as positives, with separate AUPRC computed using negatives with different non-target modifications (Ψ, m6A, m5C) and unmodified IVTs. All reference positions supported by the models were used, with all context detection available for SWARM, CHEUI ^16^, TandemMod ^15^, dorado (0.8.2), and m6ABasecaller ^22^, while DRACH-only constraint was used for m6anet ^21^ and mAFiA ^19^, and specific Ψ 5-mers were constrained by psi-co-mafia ^20^.

### Stoichiometry benchmarking

SWARM estimates stoichiometry at each tested reference site as the number of observations where read-level probability is over 0.5 out of all tested signals at a given site. To benchmark m6A stoichiometry correlation with GLORI in mRNA sites in HeLa, models were applied to publicly available HeLa nanopore samples ^58^ for SQK-RNA002, and to an in-house HeLa sample used for SQK-RNA004. Ψ and m5C mRNA stoichiometries were benchmarked against BACS ^25^ and UBS ^26^ measurements using publicly available HEK293T samples ^38^ for SQK-RNA002, and an in-house HEK293T sample for SQK-RNA004. Ψ stoichiometry in rRNA was benchmarked against mass-spec ^24^ in a publicly available HeLa *in vitro* polyadenylated sample ^58^ for SQK-RNA002, and in-house HeLa *in vitro* polyadenylated sample for SQK-RNA004. Since SingleMod SQK-RNA002 model was trained on the same HeLa samples used for benchmarks in this study, we excluded it from the m6A SQK-RNA002 stoichiometry benchmark. Pearson’s R was computed for each stoichiometry comparison, restricted only to reference sites reported by all compared methods. Transcriptomic sites aligned to the cDNA reference were lifted over to the hg38 primary assembly using R2DTool ^59^, and the stoichiometry was calculated for each genomic coordinate by averaging across covered transcripts and weighting by coverage. Minimum coverage of 20 reads per transcript and default probability cutoffs were used for all nanopore methods.

### Site-level overlap benchmarking

Nanopore-detected sites were evaluated against site labels reported by orthogonal methods using the same set of samples as described for the stoichiometry benchmarking. AUPRC was computed for each tool using site-level probabilities when available (SWARM, CHEUI, m6Anet, NanoSpa), or using stoichiometry (TandemMod, dorado 0.8.2, mAFiA, psi-co-mAfiA, m6ABasecaller). For AUPRC analysis, true modification sites were considered those reported by orthogonal methods, while negative sites were all tested sites that were not orthogonally reported. Benchmarks were restricted to reference sites tested by all compared methods. SingleMod SQK-RNA002 m6A model was excluded from the SQK-RNA002 site-level benchmarks because it was trained on the HeLa samples used in the benchmarks. Transcriptomic sites aligned to the cDNA reference were lifted over to the genome as described above, and maximal site-level probability (or stoichiometry if site-probability was not available from the method) was selected from transcripts covering the same genomic coordinate. A maximal value (probability/stoihciometry) was used across validated U homopolymers when comparing Ψ site predictions to bisulfite methods, which do not have single-base resolution within homopolymers. A minimum coverage of 20 reads per transcript site was used for all methods.

### Site-level cut-offs for *de-novo* modification detection

The false positive rate (FPR) at different site-level probability cut-offs was estimated from predictions on fully unmodified polyA+ transcriptome IVTs. Publicly available IVT samples from HeLa, A549 and HepG2 ^34,58^ were used for SQK-RNA002, while publicly available HeLa and HEK293T samples ^54^ were used for SQK-RNA004. Minimum coverage of 20 reads per transcript was used. The site-level probability was set to 0 for all sites with stoichiometry under the detection threshold (10%). FPR was calculated as the number of predicted sites with a probability larger than a given cut-off out of all tested sites in that sample. For each model, the cut-off that resulted in an FPR < 0.1% in HeLa transcriptomes was selected for *de novo* detection of modified sites in further analyses.

### Modification co-occurrence

Transcript-level m6A and Ψ co-occurrence was defined as the detection of at least one of each modification type in the same transcript. The expected number of transcripts displaying m6A and Ψ co-occurrence was computed by permuting site labels among tested sites for each tissue sample 1000 times. The mean and standard deviation of the permuted number of transcripts with co-occurrences were used to calculate z-scores as z-score =(x-μ)/σ, with x = observed value; μ = sample mean; σ = sample standard deviation. Two-tailed p-values were obtained from the z-scores using p = 2(1−Φ(∣z∣)), where Φ = the cumulative distribution function of the standard normal distribution.

Read-level co-occurrence was assessed for all m6A–Ψ site pairs detected within the same transcripts, restricted to pairs where each site was the nearest neighbour of the other modification type. For each site pair, only reads where both m6A and Ψ were tested were considered, and positions in individual reads were considered as modified if the predicted probability exceeded 0.5. The observed number of reads with both modifications detected was compared to the null distribution by permuting the tested read-level modification states within the same reference position. Permuted mean and standard deviation were used to compute z-scores and p-values using the above formula.

### Enrichment near splice-sites

m6A and Ψ sites detected across tissue samples were annotated with transcript metadata regarding the closest splice site distance and position with respect to the CDS/UTRs using R2DTool ^59^ and GTF annotation files for corresponding species (Ensembl release 109). Each transcriptomic position was considered as modified if detected above the described thresholds in at least one tissue of a given species, while background distribution was computed for transcriptomic sites tested in at least one tissue with coverage over 20 and writer motifs (DRACH for m6A, UVUAR for PUS7 Ψ, and GUUCNANNC for TRUB1 Ψ). DRACH m6A, PUS7 Ψ, and TRUB1 Ψ motifs were handled separately, and within each species, the modification site labels were shuffled 1000 times, computing modification site frequency in sliding density windows with window size=100bp, step=10bp, and considering only sites within 1kb from the closest splice junction. Observed modification frequency was compared to the permuted mean and standard deviation in each window to compute z-scores and p-values, adjusted with the Benjamini-Hochberg method.

### TRUB1 substrate secondary structure analysis

Sequence complementarity matrices were generated as in Safra et al. ^36^. RNA secondary structure of TRUB1 substrate was modeled using ViennaRNA python API version 2.7.0 on sequences 6bp upstream to 10bp downstream of the candidate pseudouridine site, i.e. NNNNGU**U**CNANNCNNNN, using `(((((.)))))` as a constraint.

### Nascent RNA pseudouridine detection at TRUB1-candidate sites

Raw nanopore signals from Sethi et al. ^44^ were basecalled with dorado 0.7.2 and aligned to the hg38 primary assembly using minimap2 -ax splice. Reads covering the region of each target site were extracted using samtools view and pod5 subset. Target reads were basecalled with dorado 0.8.2 and pseU model, converted to fastq with -T* flag and aligned to the genome using minimap2 -ax splice -y to keep modification tags for dorado analysis. To perform reference-free SWARM modification detection in target reads, dorado 0.7.2 basecalls were aligned using minimap2 -ax map-ont and resquiggled via f5c eventalign using a reference with basecalled read sequences, followed by SWARM read-level modification detection. SWARM modsam predictions were converted to fastq and aligned to the genome with same commands as described for dorado 0.8.2. Aligned mod.bam files were processed to only contain modification readouts at target sites, and modification probabilities over 0.9 were considered as modified for IGV visualisation and stoichiometry quantification. Minimum coverage of 5 reads was used for stoichiometry quantification.

### *In vitro* TRUB1 pseudouridylation nanopore analysis

Raw nanopore data was basecalled using dorado 0.7.2 for SWARM analysis and dorado 0.8.2 with pseU model for Ψ detection. Basecalled reads were mapped to the barcoded reference constructs (Supp. Table 7) using minimap 2.24 -ax map-ont –y. Reads were filtered for primary alignments on the positive strand using samtools view –F 2324. For SWARM analysis, mapped reads were resquiggled using f5c eventalign with same flags as in methods above with additional --min-recalib-events 50 to handle tRNA reads. Modification stoichiometry was calculated stringently using >0.9 cutoff for modified calls and <0.5 for unmodified calls.

## Data availability

The specific use of each nanopore sample for training, testing, or biological discovery is detailed in Supp. Table 8, along with the orthogonal datasets used for training and testing. Datasets produced for this work are uploaded to the ENA project PRJEB106788. These include synthetic IVT-i and IVT-m samples (SQK-RNA002 & SQK-RNA004), unmodified IVT RNA from human hippocampus and mouse embryo cDNA libraries (SQK-RNA004), HeLa size-selected *in vitro* polyadenylated (SQK-RNA004), HEK293T WT (SQK-RNA004), and TRUB1-incubated synthetic isoform IVT samples (SQK-RNA004). Curlcake modified IVTs (SQK-RNA002) were obtained from SRA projects PRJNA563591 and PRJNA511582. Datasets from mammalian tissues (SQK-RNA002) ^7^ were obtained from SRA (PRJNA1188790). Datasets for unmodified cell-line transcriptomes (HeLa, A549, HepG2) (SQK-RNA002) and HeLa WT mRNA and rRNA (SQK-RNA002) ^34,58^ were obtained from SRA (PRJNA947135 and PRJNA777450). Datasets for HEK293T WT and METTL3-KO (SQK-RNA002) ^38^ were obtained from ENA (PRJEB40872). Datasets for HEK293T WT and TRUB1-KD (SQK-RNA002) were obtained from ENA (PRJEB72637). Datasets for HeLa WT and PUS7-KD (SQK-RNA004) were obtained from GEO (GSE314420). Datasets for HEK293T IVT and HeLa IVT (SQK-RNA004) ^54^ were obtained from ENA (PRJEB80229). Datasets for HeLa WT mRNA and pre-mRNA (SQK-RNA004) ^44^ were obtained from SRA (PRJNA1256087).

## Software Availability

SWARM v1.0 (https://github.com/comprna/SWARM)

Minimap2 (https://github.com/lh3/minimap2/releases/tag/v2.24)

Dorado 0.7.2 & 0.8.2 (https://github.com/nanoporetech/dorado/releases/)

TandemMod v1.1.0 (https://github.com/yulab2021/TandemMod)

M6Abasecalller (https://github.com/novoalab/m6ABasecaller)

CHEUI v1.0 (https://github.com/comprna/CHEUI)

NanoSpa (https://github.com/sihaohuanguc/NanoSPA)

Psi-co-mafia (https://github.com/dieterich-lab/psi-co-mAFiA)

M6Anet v2.1.0 (https://github.com/GoekeLab/m6anet)

SingleMod v1.0 (https://github.com/xieyy46/SingleMod-v1)

F5c v1.5 (https://github.com/hasindu2008/f5c/releases/tag/v1.5)

Battery-eel v0.7.2 (https://github.com/Psy-Fer/buttery-eel/releases)

Slow5tools v.1.1.0 (https://github.com/hasindu2008/slow5tools/releases/tag/v1.1.0)

R2DTool v2.0.1 (https://github.com/comprna/R2Dtool/releases/tag/v2.0.1)

ViennaRNA v2.7.0 (https://github.com/ViennaRNA/ViennaRNA/releases/tag/v2.7.0)

Tensorflow v2.15.0 (https://github.com/tensorflow/tensorflow/releases/tag/v2.15.0)

Scikit-learn v.1.4.0 (https://github.com/scikit-learn/scikit-learn/releases/tag/1.4.0-1)

Pysam v.0.22.1 (https://github.com/pysam-developers/pysam/releases/tag/v0.22.1)

Samtools v.1.22.1 (https://github.com/samtools/samtools/releases/tag/1.22.1)

## Acknowledgements

We are grateful to the personnel from the Biomolecular Resource Facility at JCSMR (ANU) for their technical assistance. We are also grateful to the personnel of the Ecogenomics and Bioinformatics Lab, a joint initiative of the Research School of Biology (ANU) and the Commonwealth Scientific and Industrial Research Organisation (CSIRO), and particularly to Niccy Aitken and Ashley Jones, for their continued support and feedback regarding ONT sequencing.

