## Supplementary Figures for "SWARM resolves nanopore signal interference between RNA modification types and reveals splicing-shaped pseudouridylation"

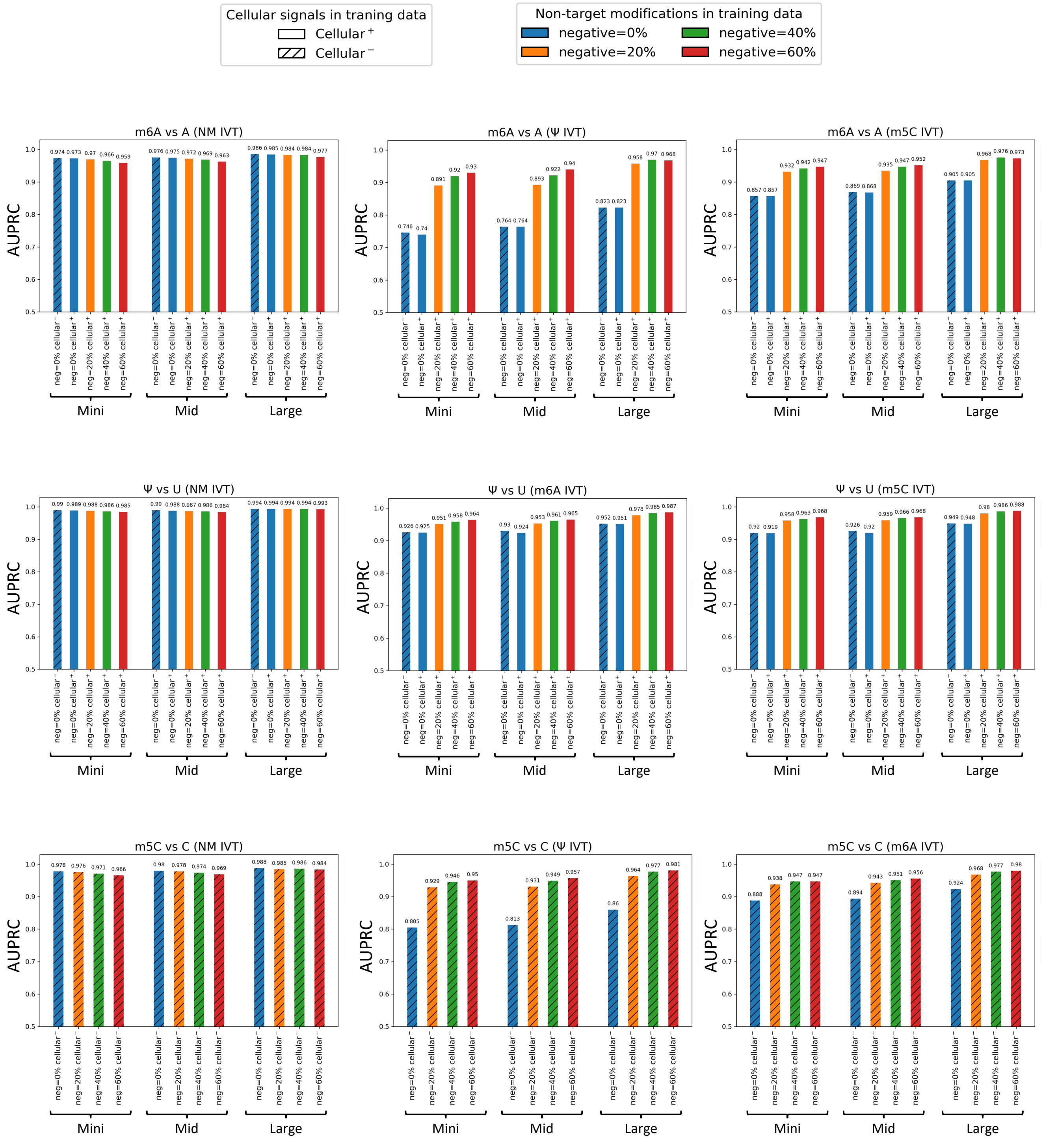


**Supplementary Figure 1. Evaluation of training datasets for read-level modification specificity.** Area under the Precision-Recall curves (AUPRC) (y-axis) values are shown in bar plots for all read-level models trained with specified training datasets and network architectures (Mini/Mid/Large). Percentage of negative training data containing signals from non-target modification IVT samples is shown with colours indicated in the legend, i.e. negative=20% for m6A model means that 20% of negative signals are A sites from Ψ, m5C, and ac4C IVT samples, and remaining 80% percent is from non-modified IVT. Inclusion of cellular data during training is indicated with bar hatches, with clear bars labelled as cellular$^+$ containing cellular data, and hatched bars labelled as cellular$^-$ not trained with cellular signals. Title of each panel shows the target modification for each model, and sample used for negative data for computing Precision-recall AUC, i.e. m6A vs A (Ψ IVT) means that m6A model was evaluated using A sites from m6A IVT as positives and A sites from Ψ IVT as negatives. Non-modified IVT samples are abbreviated as NM. Evaluation was conducted on SQK-RNA002 data.

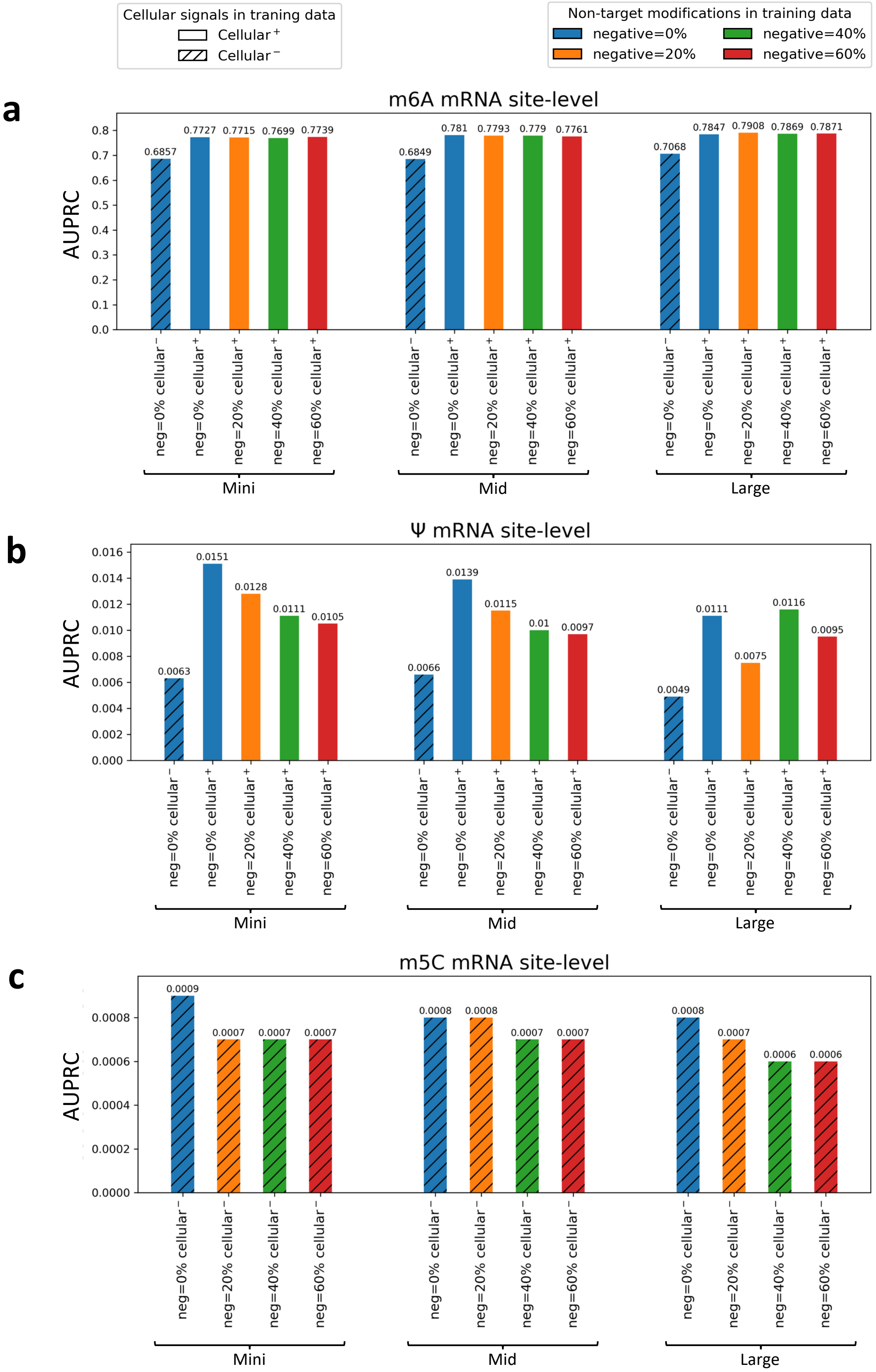


**Supplementary Figure 2. Evaluation of training datasets for site-level modification detection in cell lines.** Each panel shows mRNA site-level Area Under the Precision-Recall Curves (AUPRC) evaluated with orthogonal datasets using a site-level network with same (Mini) architecture trained for each read-level model with specified training data compositions. Training datasets differed in inclusion of cellular data in training (cellular$^{+}$ or cellular$^{-}$), and amount of non-target negative modifications with shown percentages, outlined in more detail in description of Supp. Fig. 1. Each of the three panel columns shows a read-level network architecture, Mini on left, Mid in the middle, and Large on right. For m6A **(a)**, we used GLORI on Hek293T DRACH data, for Ψ **(b)**, BACS on HeLa data, for m5C **(c)**, UBS on HeLa data. Evaluation was conducted on SQK-RNA002 data.

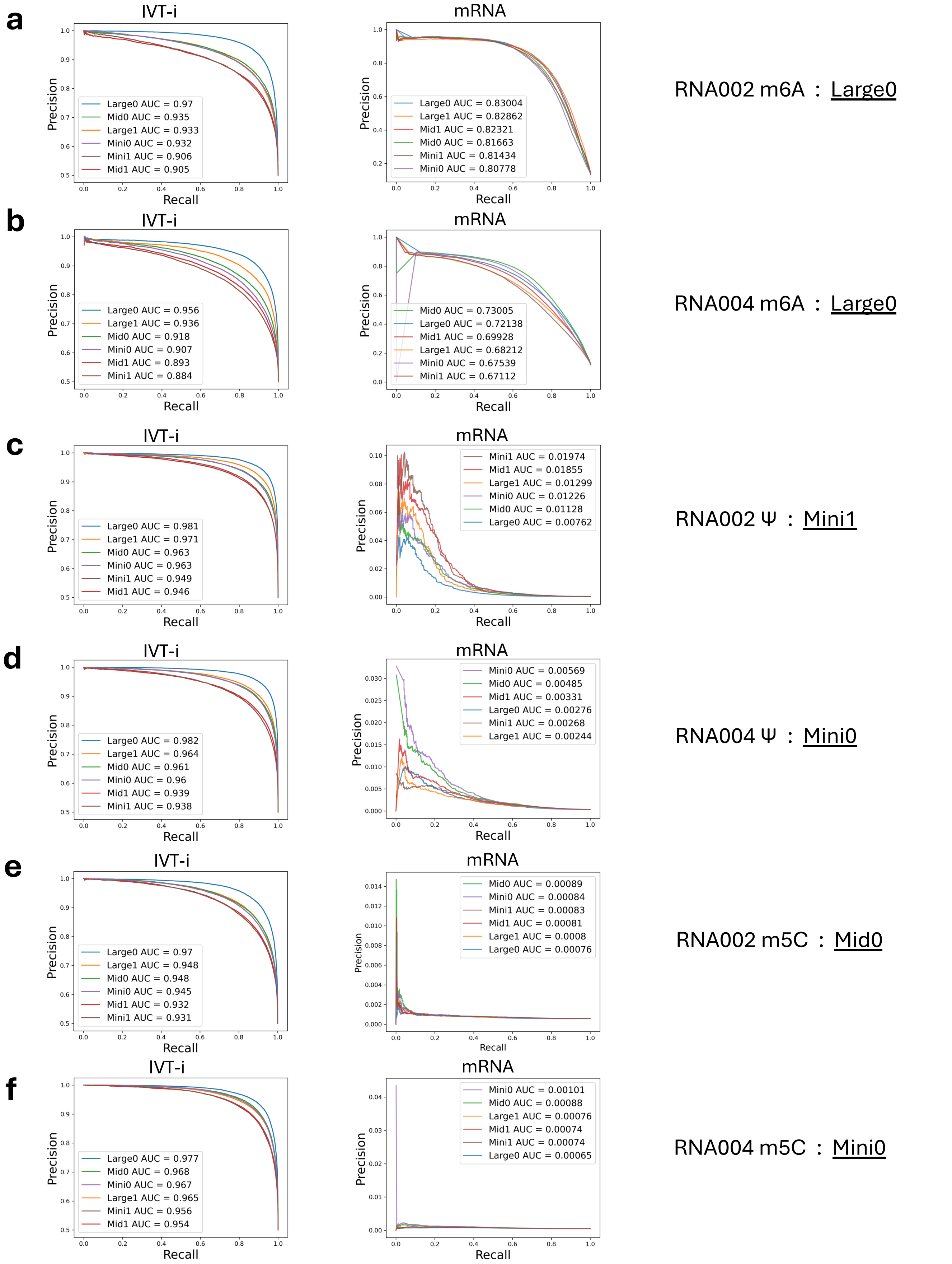


**Supplementary Figure 3. Selection of read-level model hyperparameters.** Precision-recall curves are shown for different read-level models across both ONT chemistries, for m6A **(a,b)**, Ψ **(c,d)**, and m5C **(e,f)**. Two metrics were used: left panels show read-level precision-recall curves on the IVT-i test split signals; the right panels show mRNA site-level precision-recall curves evaluated with orthogonal datasets using a site-level network with same (Mini) architecture trained for each read-level model. For m6A (a,b), we used GLORI on Hek293T data, for Ψ (c,d), BACS on HeLa data, for m5C (e,f), UBS on HeLa data. Three different model architectures were tested (Mini, Mid, Large) trained with (1) or without (0) sample weights. Selected models are shown in text to the right.

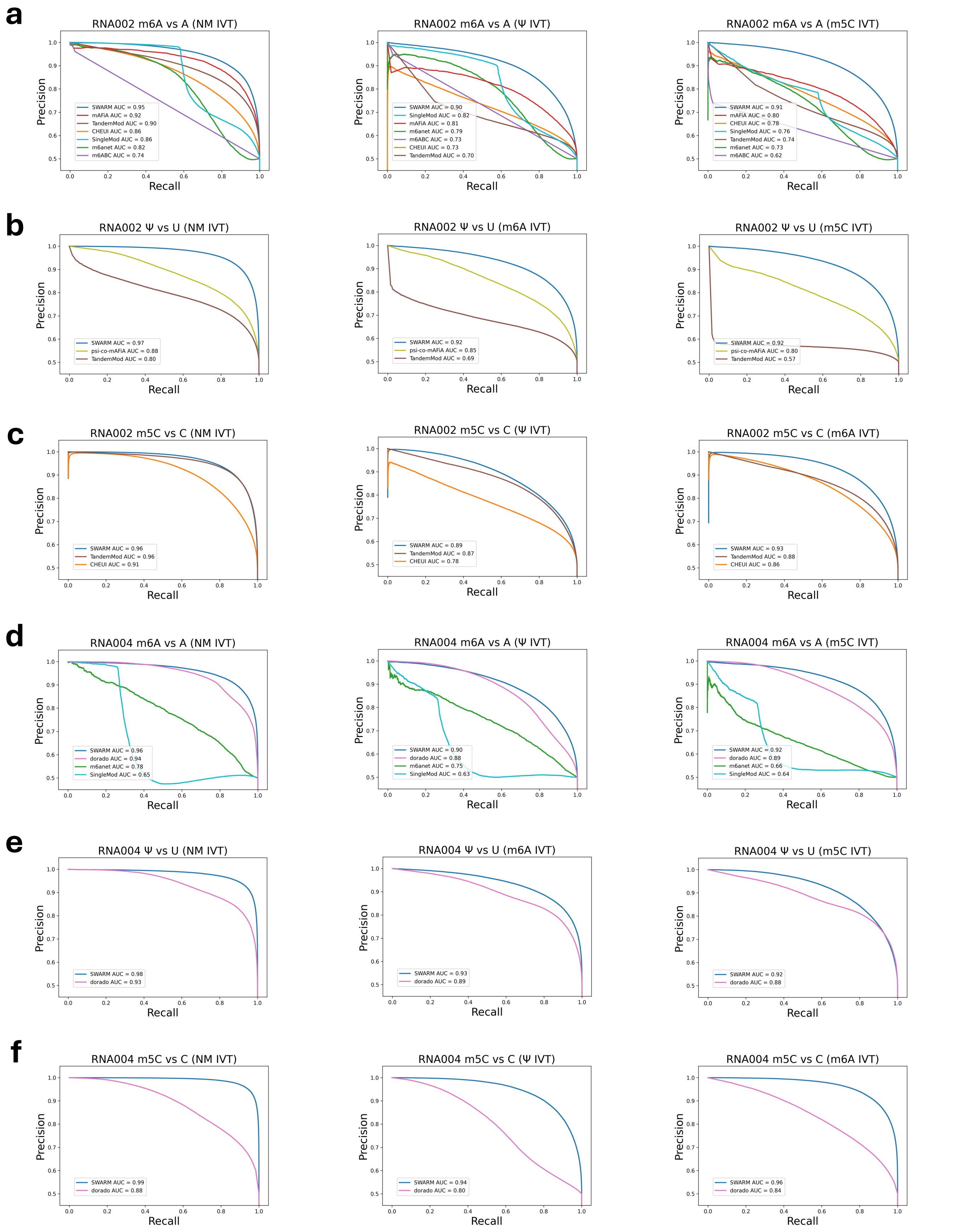


**Supplementary Figure 4. Precision-recall curves for the single-read accuracy analysis.** The plots display the precision-recall curves calculated for read-level modification tools for both ONT chemistries, on IVT M1 and M2 for SQK-RNA002 **(a-c)** and on IVT M4 for SQK-RNA004 **(d-f)**, at single-nucleotide resolution. Benchmarking datasets were built from independent synthetic IVTs not used for training. Positive cases were built from reads containing the modification. Various negative sets were considered, encompassing reads without modifications (NM) or reads with non-target modifications.

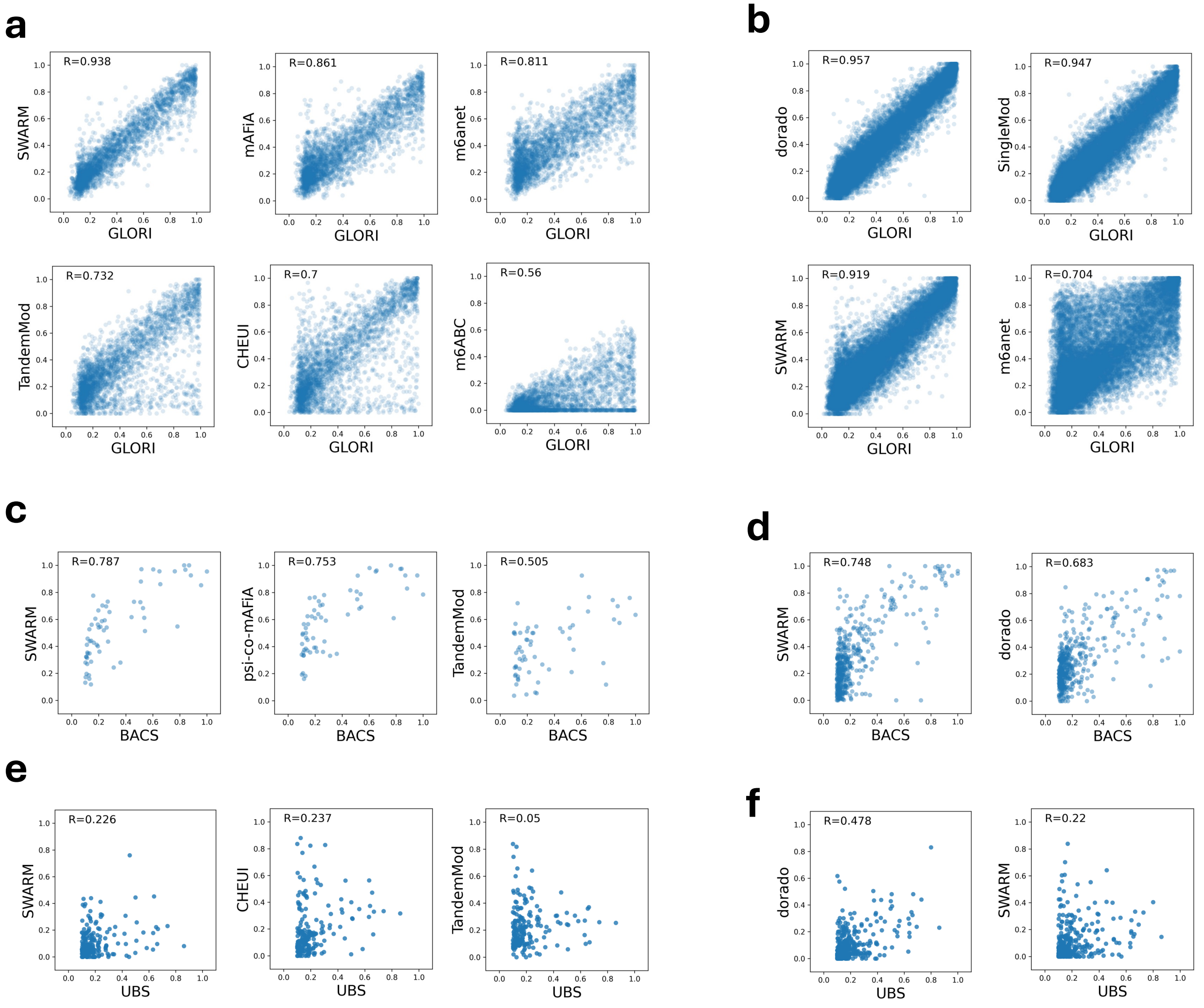


**Supplementary Figure 5. Correlation of stoichiometries with orthogonal methods.** The X-Y plots show the stoichiometry correlations on mRNA sites predicted by various methods (y-axis) for m6A in HeLa in SQK-RNA002 **(a)** and SQK-RNA004 **(b)** compared with GLORI (x-axis), for Ψ in HeLa in SQK-RNA002 **(c)** and SQK-RNA004 **(d)** compared with BACS (x-axis), and for m5C in Hek293T in SQK-RNA002 **(e)** and SQK-RNA004 **(f)** compared with UBS (x-axis). The correlation values are shown in Figure 2.

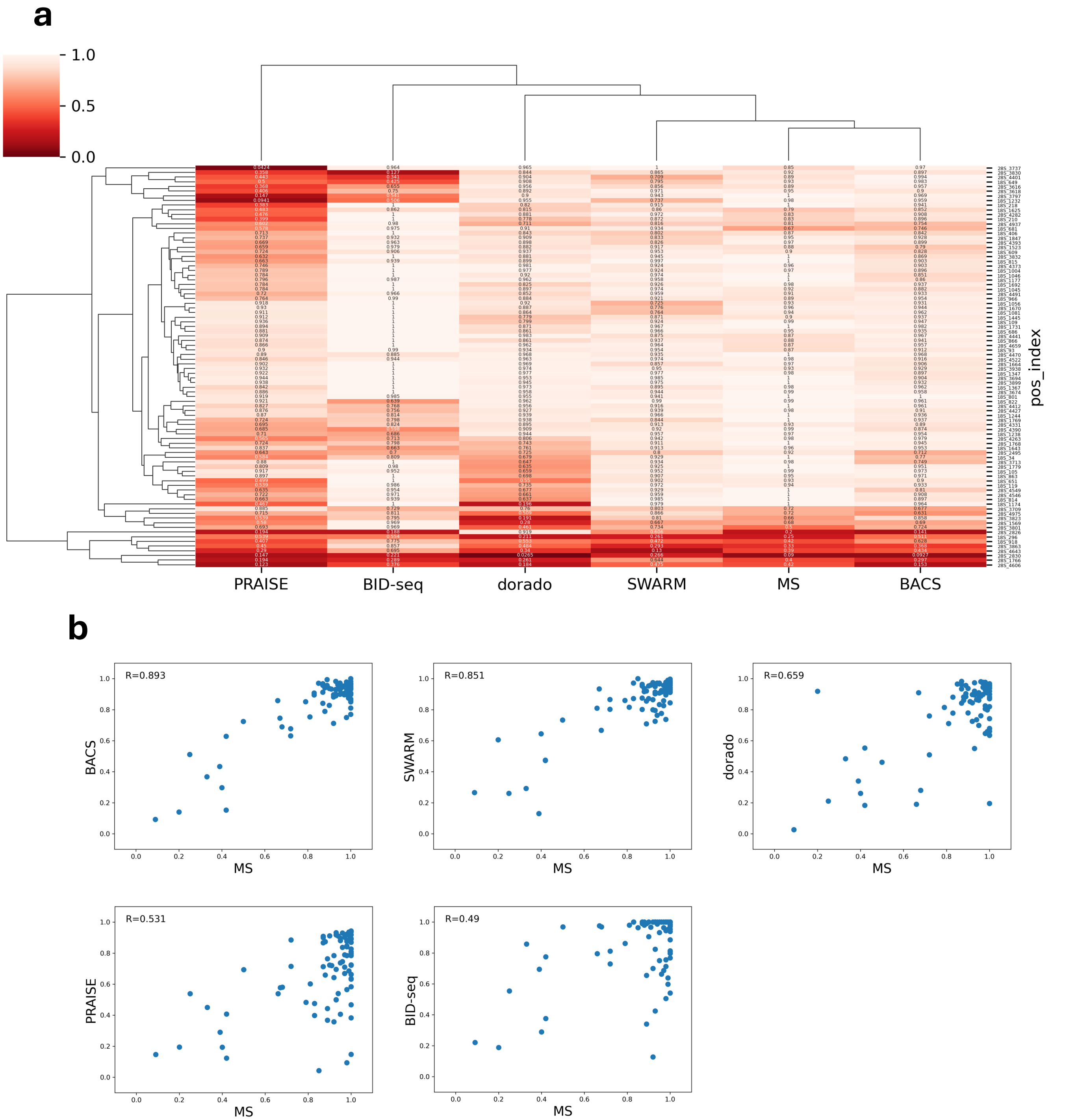


**Supplementary Figure 6. Pseudouridine stoichiometries in HeLa rRNA across orthogonal methods. (a)** Hierarchical clustering of the stoichiometry values (ranking from 0 to 1) estimated by each method (x-axis) on rRNA pseudouridine sites. ONT SQK-RNA004 DRS data from HeLa cells. **(b)** Pairwise comparison of the stoichiometry values estimated by each method in (a).

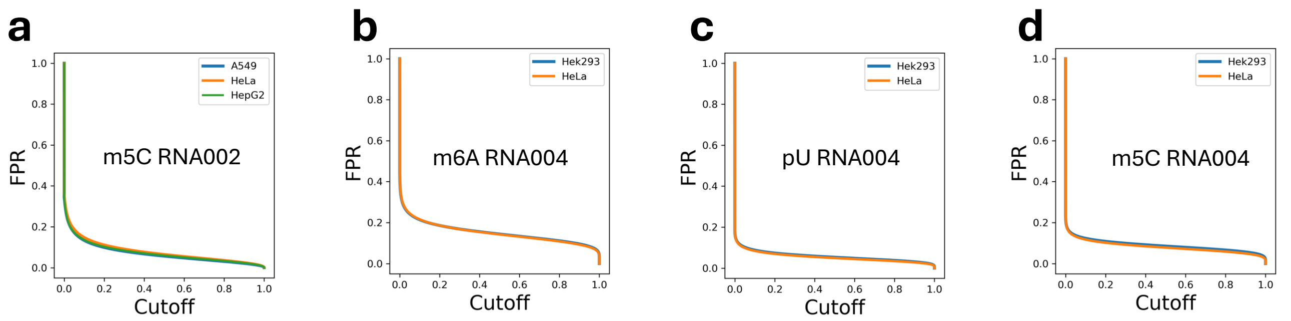


**Supplementary Figure 7. False positive rate (FPR) on fully unmodified in-vitro transcribed human cell lines.** **(a)** FPR for the SWARM m5C RNA002 model on A549 (blue), HeLa (orange), and HepG2 (green) cell lines. **(b-d)** FPR for the SWARM RNA004 m6A, pU, and m5C models on Hek293T (blue) and HeLa (orange) cell lines.

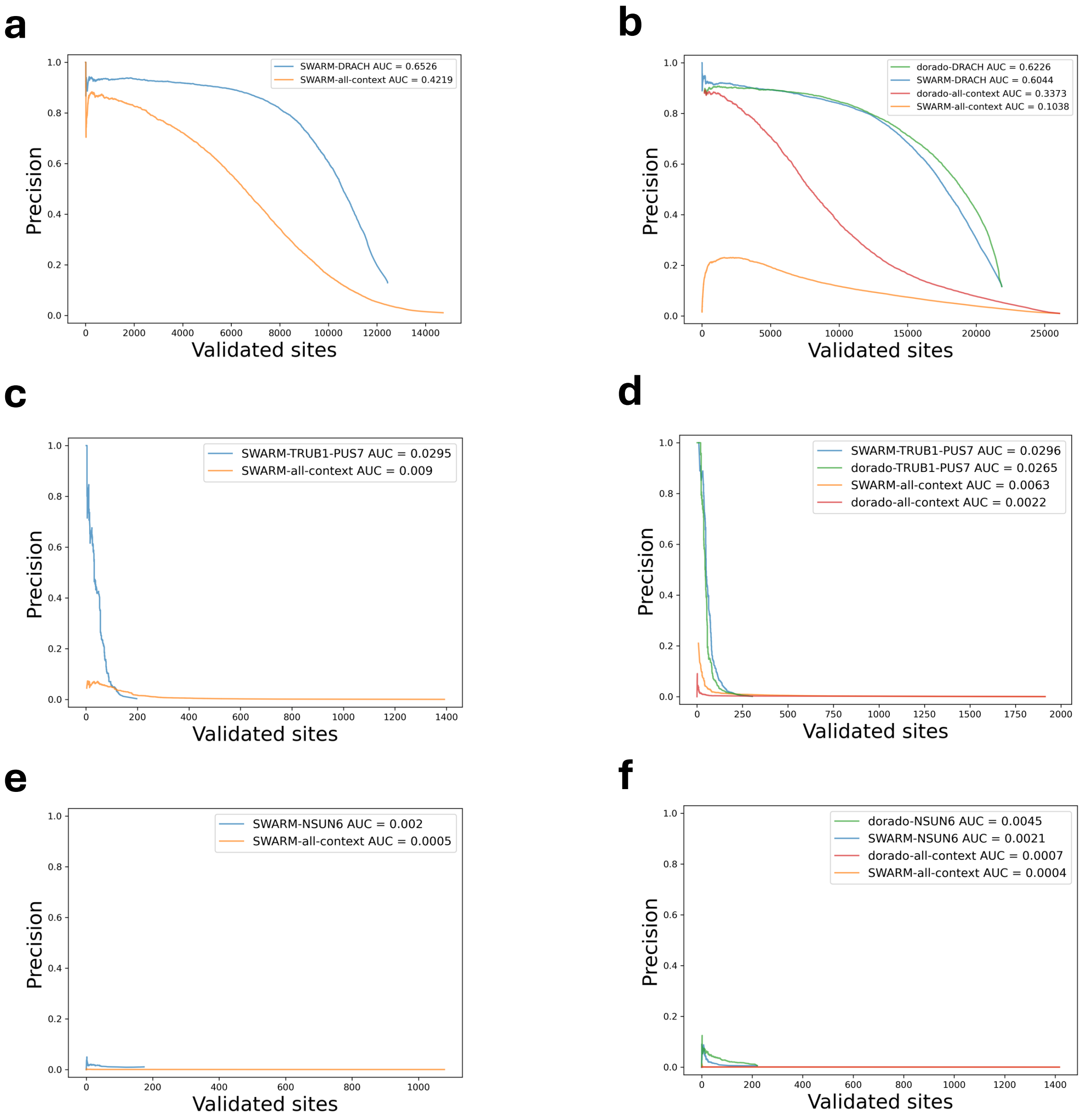


**Supplementary Figure 8. Precision evaluation of site-level predictions in all-context vs biologically-relevant motifs.** Precision (y-axis) of SWARM's site-level predictions as a function of the number of validated sites detected (x-axis). Validated sites were considered to be those detected by GLORI for m6A **(a, b)**, BID-seq and PRAISE for Ψ **(c, d)**, and UBS-seq for m5C **(e, f)**. In each case, we analysed SQK-RNA002 (left panels) and SQK-RNA004 data (right panels). We considered the predictions in all sequence contexts, or restricting to the expected motifs: DRACH for m6A, GUΨCNA (TRUB1) and UNΨAR (PUS7) for Ψ, and CUCCA (NSUN6) sites.

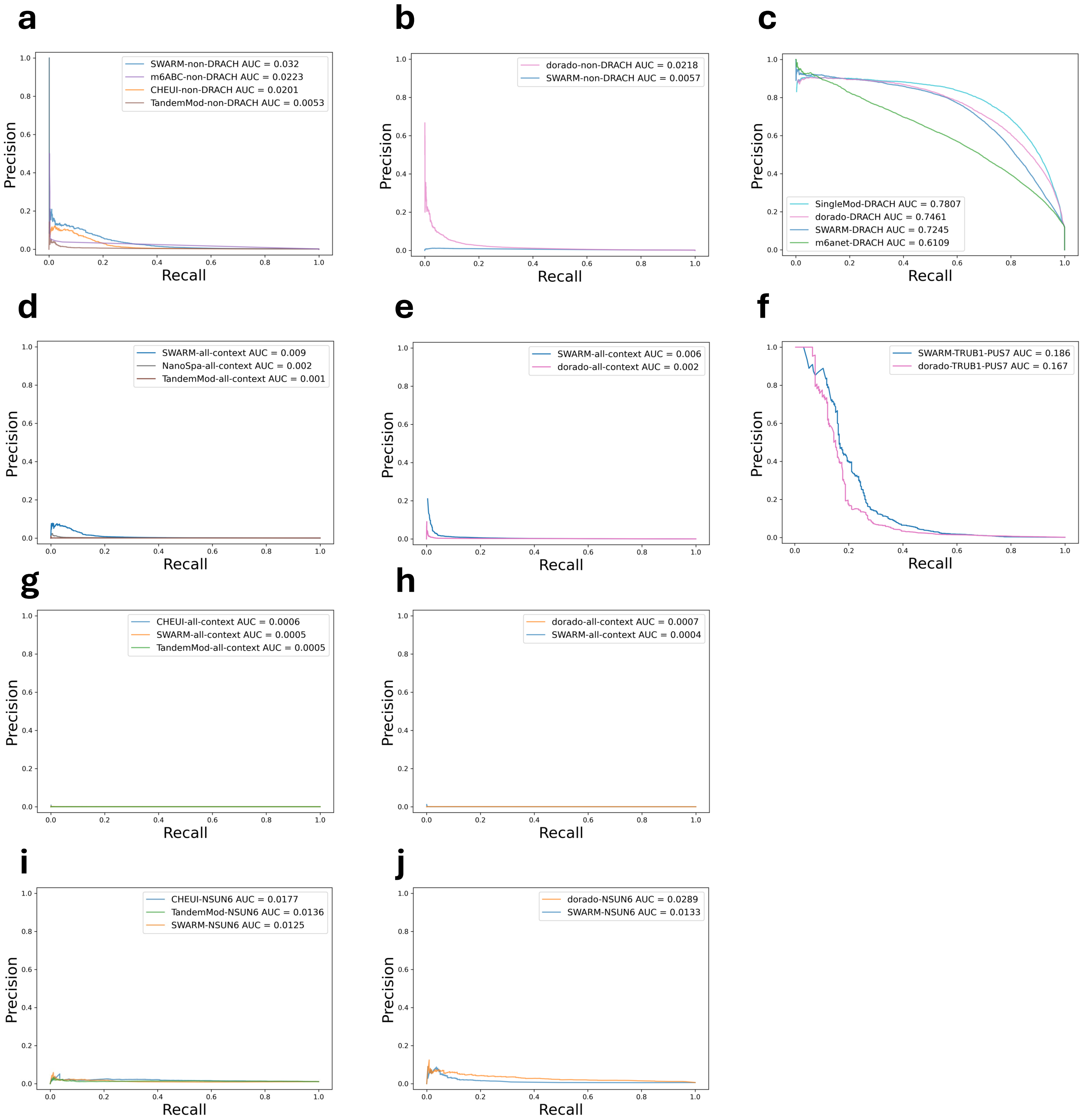


**Supplementary Figure 9. Precision-recall evaluation of site-level predictions on cellular mRNA.** Precision (y-axis) versus recall (x-axis) curves of SWARM's site-level predictions. Validated sites were considered to be those detected by GLORI for m6A **(a-c)**, BID-seq and PRAISE for Ψ **(d-f)**, and UBS-seq for m5C **(g-j)**.

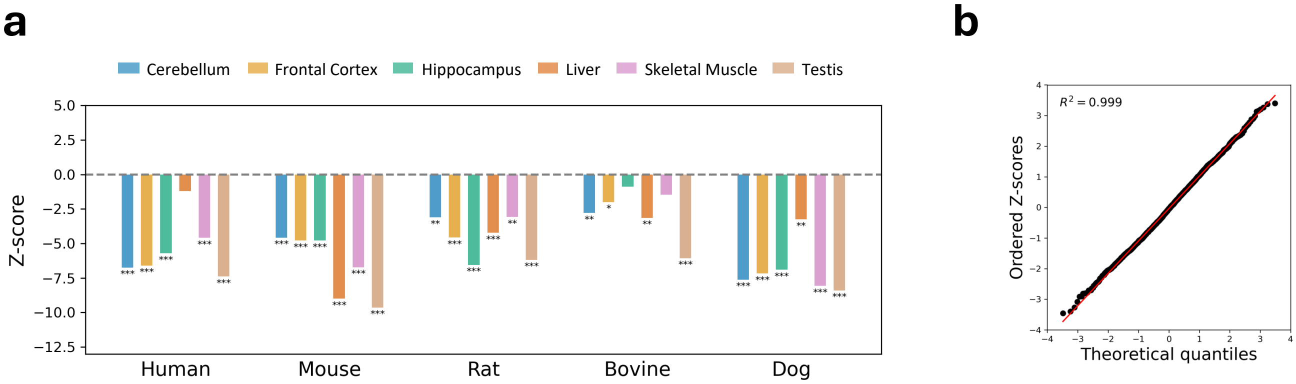


**Supplementary Figure 10. Analysis of m6A and pseudouridine co-occurrence. (a)** Z-scores of the comparison of observed co-occurrences at the transcript level with the values obtained after permuting 1000 times all sites tested. **(b)** QQ-plot of the observed z-scores (y-axis) and theoretical quantiles (x-axis) for the analysis of the read-level co-occurrence of m6A, at DRACH sites, and Ψ, at TRUB1 (GUΨCNANNC) or PUS7 (UNΨAR) sites.

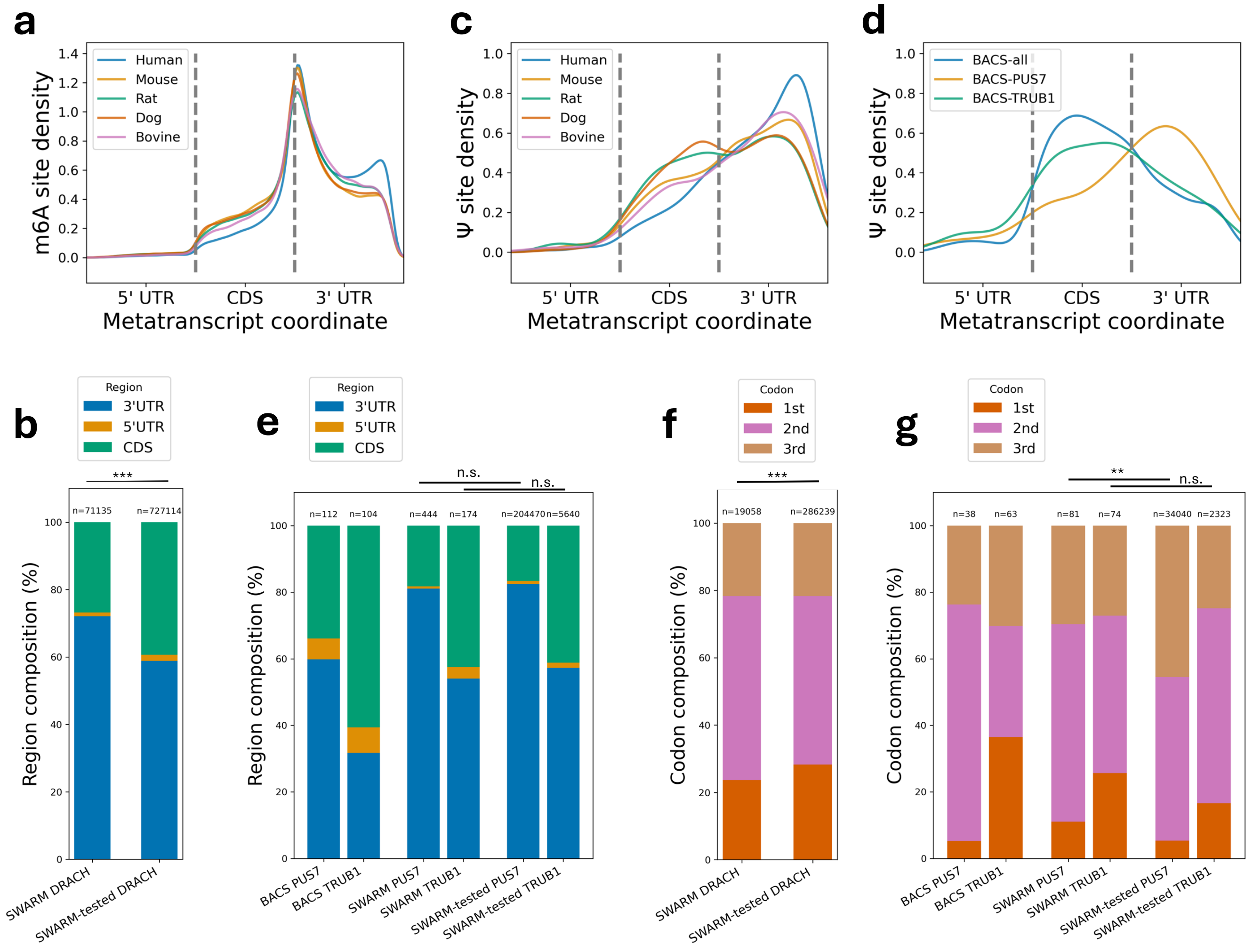


**Supplementary Figure 11. Modification distribution with respect to transcript elements. (a)** Density plot of m6A sites detected by SWARM in at least one tissue of each species with respect to the location of UTR and CDS elements. **(b)** Composition of m6A sites detected by SWARM in human tissues broken down by UTR and CDS regions, comparing detected sites (left) and tested sites (right) with candidate DRACH motifs. **(c-d)** Density plot of Ψ sites detected by SWARM in at least one tissue of each species (c) and BACS sites in HeLa (d) with positions mapped to the UTR and CDS elements. **(e)** Region composition of UTR and CDS elements within Ψ sites detected by BACS in HeLa (left pair) and detected by SWARM (middle pair) or tested by SWARM (right pair) in human tissues. Each bar pair is split by Ψ writer motifs, with PUS7 in left and TRUB1 in right. **(f)** Breakdown of codon positions of SWARM detected (left) and tested (right) m6A DRACH sites in human tissues. **(g)** Percent of Ψ sites in each codon position for BACS sites in HeLa (left pair), detected by SWARM (middle pair) or tested by SWARM (right pair) in human tissues. Bar pairs are split by Ψ writer motifs, PUS7 is left and TRUB1 is right. Statistical significance of chi-squared test is shown above bars: n.s. = non-significant; * = 0.05; ** = 0.005; *** = 0.001.

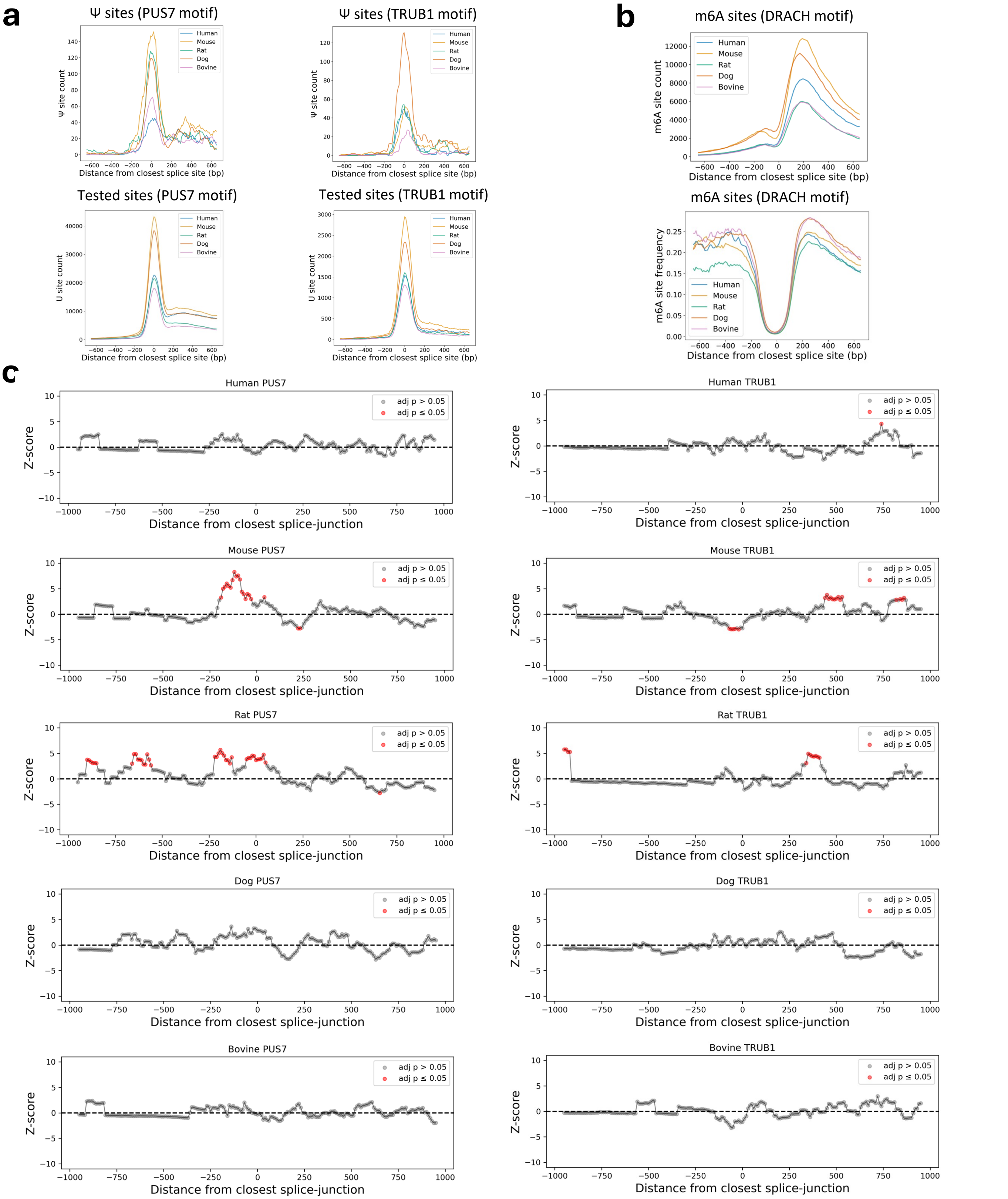


**Supplementary Figure 12. Analysis of modification enrichment across exon-exon boundaries. (a)** Counts of Ψ detected in at least one tissue (upper panels) and tested (lower panels) U sites with PUS7 (UVΨAR, V=A,C,G) and TRUB1 (GUΨCNANNC) motif contexts in 100bp windows centred on the closest splice site. **(b)** Counts (upper panel) and frequency (lower panel) of m6A sites detected in DRACH motifs in at least one tissue of tested species in 100bp windows centred on the closest splice site. **(c)** Enrichment of detected Ψ sites at different distances from the closest splice site given the distribution of the tested sites in each species with PUS7 (left panels) and TRUB1 (right panels) motifs. Z-scores obtained using the observed counts of detected Ψ in 100bp windows against 1000 permutations of the tested sites in the same windows.

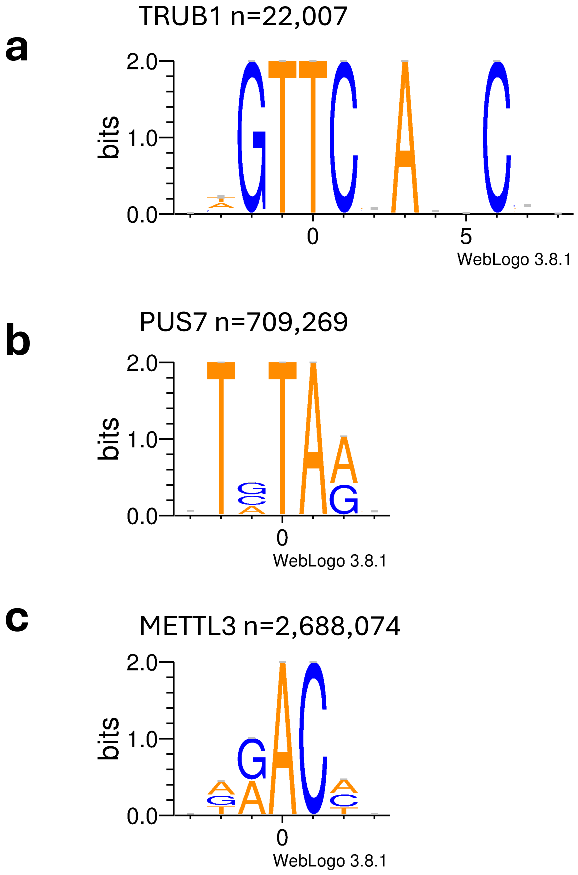


**Supplementary Figure 13. Motifs of non-predicted candidate Ψ sites.** Motifs of candidate TRUB1 **(a)**, PUS7 **(b)**, and METTL3 **(c)** sites with no evidence of modification in any tissue across five mammals. Sites were listed as unmodified if stoichiometry was measured under 10% in every tested tissue of given species.

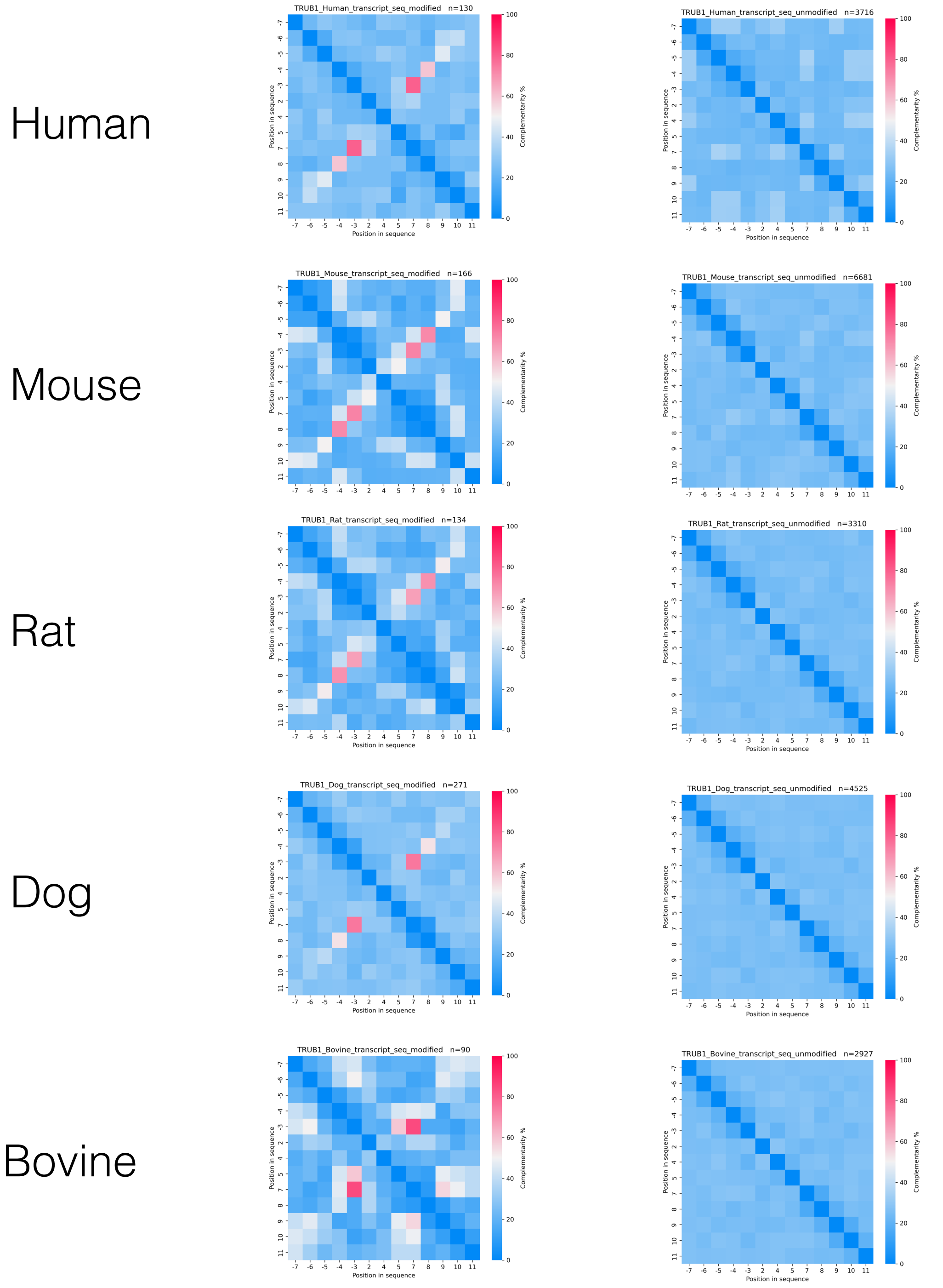


**Supplementary Figure 14. Sequence complementary of Ψ sites with TRUB1 motifs.** Complementarity of all TRUB1 sites in each species calculated using the same method from Safra et al. 2017.

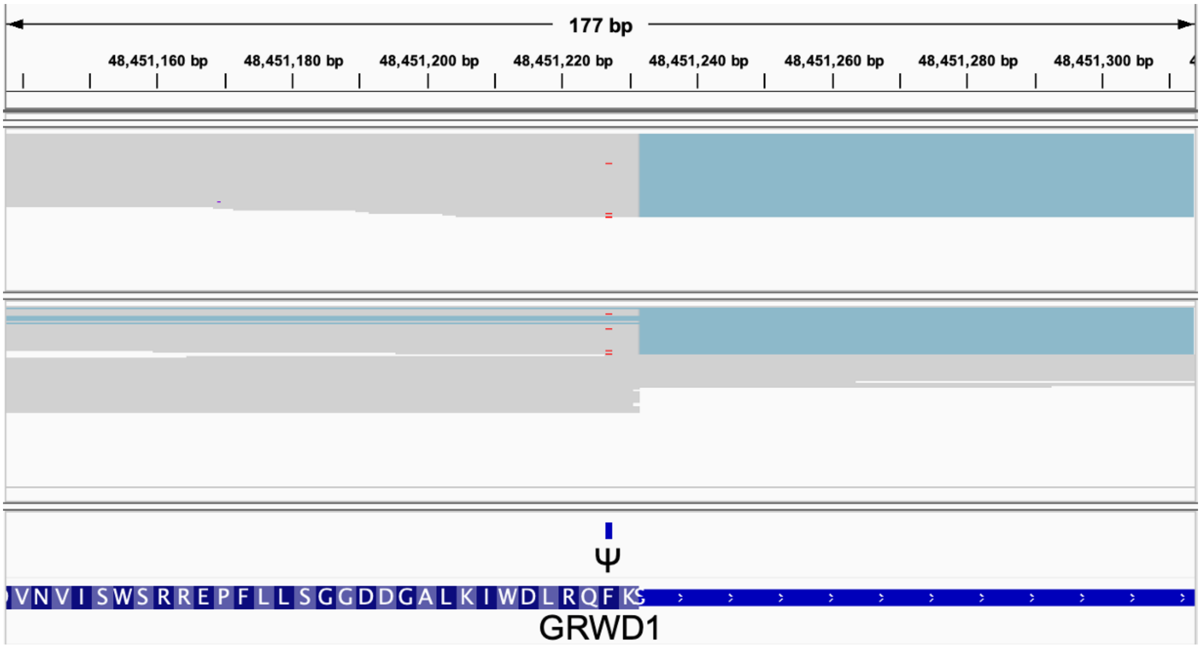


**Supplementary Figure 15. Validation of splicing-dependent TRUB1 pseudouridine sites with pre-mRNA direct RNA sequencing.** IGV plot showing in red the SWARM-predicted Ψ sites in individual reads for the human gene GRWD1, obtained from the direct sequencing of pre-mRNA (Sethi et al. 2025).

| | Total untreated | | Nascent untreated | | | | Nascent PlaB+ | | | |
|---|---|---|---|---|---|---|---|---|---|---|
| | spliced | | spliced | | unspliced | | spliced | | unspliced | |
| | SWARM | dorado | SWARM | dorado | SWARM | dorado | SWARM | dorado | SWARM | dorado |
| 2:27326828 | 83 | 100 | 86 | 74 | 83 | 75 | 88 | 85 | 90 | 82 |
| 1:33014553 | 81 | 81 | 38 | 56 | 0 | 0 | 76 | 83 | 6 | 9 |
| 12:53443441 | 70 | 85 | 82 | 93 | 24 | 17 | 74 | 94 | 10 | 41 |
| 1:35603333 | 67 | 69 | 50 | 62 | 0 | 5 | 60 | 82 | 10 | 13 |
| X:154765961 | 55 | 54 | 19 | 17 | 4 | 0 | 44 | 54 | 23 | 18 |
| 17:40295391 | 38 | 50 | 27 | 35 | 10 | 10 | 50 | 52 | 9 | 11 |
| 10:3112272 | 33 | 33 | 3 | 3 | 0 | 0 | 14 | 13 | 0 | 0 |
| 8:116863212 | 29 | 30 | 23 | 14 | 0 | 0 | 33 | 38 | 0 | 0 |
| 1:215608091 | 26 | 36 | 19 | 20 | 0 | 0 | 24 | 25 | 0 | 22 |
| 22:19177964 | 13 | 23 | 14 | 0 | 0 | 2 | 12 | 37 | 0 | 0 |

Ψ stoichiometry (%): 0 – 20 – 40 – 60 – 80 – 100

**Supplementary Figure 16. Pseudouridine stoichiometry of sites with single-exon TRUB1 motifs in pre-mRNA direct RNA sequencing.** Heatmap shows Ψ stoichiometry measured by SWARM and dorado in total and nascent RNA fractions (Sethi et al. 2025) with or without PlaB treatment, and in reads with spliced/unspliced status of the closest intron. All sites have the TRUB1 substrate motif fully contained within one exon.

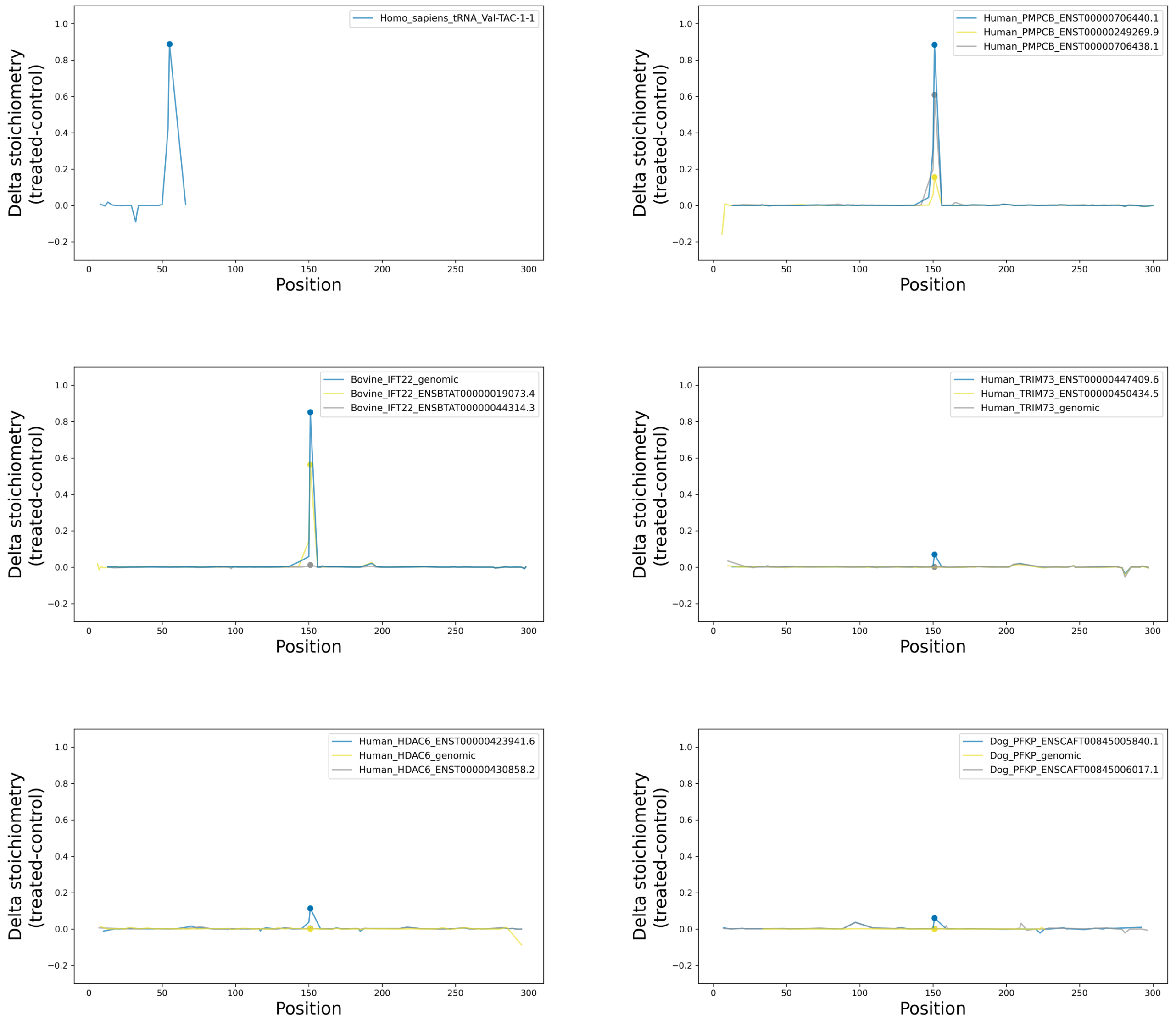


**Supplementary Figure 17. *In vitro* stoichiometry changes after TRUB1 treatment in tested genes.** Each panel shows normalised Ψ stoichiometry (TRUB1 treated - control) across tested isoforms in the given gene. Nanopore Ψ stoichiometry was averaged between SWARM and dorado at each position within the same condition. Different isoform constructs are indicated with colours in the legend, with blue isoform having the highest expected affinity to TRUB1, while yellow and grey were expected to show lower affinity. Candidate TRUB1 sites expected to be modified based of the canonical GU**U**CNANNC motif are highlighted with dotted markers.

ENST00000249269.9 (canonical) | ENST00000706438.1 (intron-retained = pre-mRNA) | ENST00000706440.1 (exon-skipping)

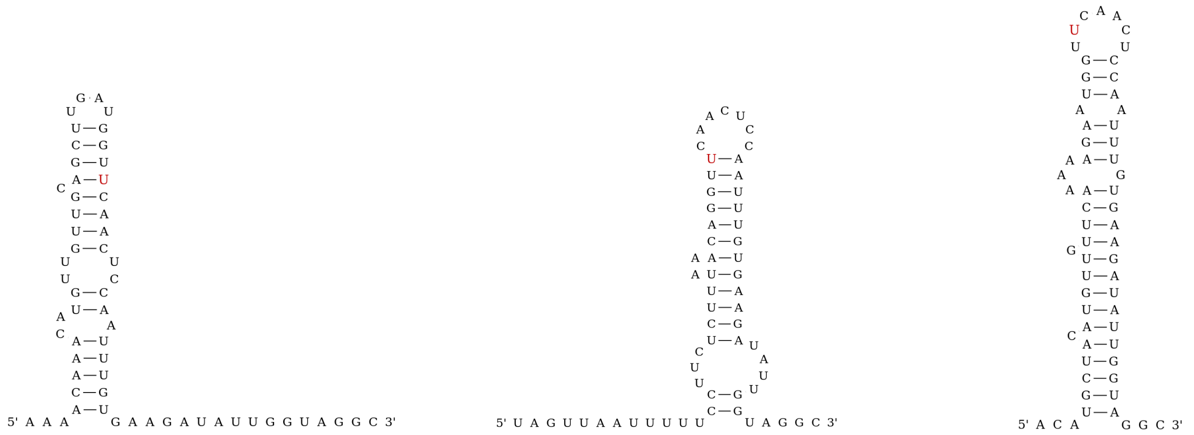

**Supplementary Figure 18. Structure of different PMPCB isoforms around candidate TRUB1 sites tested *in vitro*.** RNA structures were predicted using RNAfold from the Vienna package using 30 nucleotides upstream and downstream from the candidate TRUB1 sites, indicated in red.
